# The molecular mechanism and activity of *Kuenenia stuttgartiensis* hydrazine synthase

**DOI:** 10.64898/2026.02.25.707902

**Authors:** Femke J. Vermeir, Suzanne C. M. Haaijer-Vroomen, Pieter M. M. van der Velden, Rob Mesman, Robert S. Jansen, Wouter Versantvoort, Laura van Niftrik

## Abstract

Anaerobic ammonium-oxidizing (anammox) bacteria convert ammonium and nitrite into dinitrogen gas via the intermediates nitric oxide and hydrazine. To produce hydrazine, anammox bacteria harbor a biochemically unique enzyme: hydrazine synthase. Based on the hydrazine synthase crystal structure it was hypothesized that hydrazine is produced in a two-step mechanism. In this hypothesis, nitric oxide is first reduced to hydroxylamine (first half-reaction), followed by condensation of hydroxylamine with ammonium to hydrazine (second half-reaction). Here, we experimentally investigated the proposed molecular mechanism of hydrazine synthase and characterized and optimized the *in vitro* activity. First, we optimized the activity of isolated hydrazine synthase from anammox bacterium *Kuenenia stuttgartiensis* strain MBR1 via an anaerobic isolation method. We further compared hydrazine synthase activity measured via a coupled assay versus that of a newly established direct LC-MS assay. Next, the hypothesized second half-reaction was investigated via the direct LC-MS assay, quantifying biologically produced hydrazine from hydroxylamine and ammonium. Despite variation in hydrazine synthase activity across assays, we determined ammonium and hydroxylamine affinity and investigated product inhibition. Finally, we found that hydrazine synthesis from ammonium and hydroxylamine by isolated hydrazine synthase is oxygen-tolerant, strongly suggesting that the second half-reaction is initiated on an oxidized heme within hydrazine synthase. Taken together, the results corroborate that condensation of ammonium with hydroxylamine to form hydrazine is the second half-reaction of the proposed two-step mechanism for hydrazine synthesis by hydrazine synthase.

## Introduction

Anaerobic ammonium oxidation (anammox) is the anaerobic microbial process in which ammonium oxidation is coupled to nitrite reduction to produce dinitrogen gas. To date, anammox bacteria are the only microorganisms known to perform the anammox reaction. Anammox bacteria are ubiquitous in nature and are found in oxygen minimum zones, marine sediments, freshwater lakes and multiple other oxygen-limited natural environments (Oshiki et al., 2016b). Due to their abundance and metabolism, anammox bacteria significantly contribute to the removal of fixed nitrogen from the environment (Devol, 2015; Lotti et al., 2015) and can be applied in wastewater treatment (Lackner et al., 2014).

In the laboratory, anammox bacteria are cultivated in bioreactors with up to 95% enrichment, where they generally grow in larger cell aggregates with doubling times varying from days to weeks (Lotti et al., 2015; Zhang et al., 2017). The molecular mechanism and physiology of the anammox reaction was, and is, studied in the model anammox species “*Candidatus* Kuenenia stuttgartiensis”, that can be cultivated in planktonic enrichment cultures (van der Star et al., 2008). The catabolic anammox reaction takes place in a dedicated compartment termed the anammoxosome. Anammox proceeds via nitrite reduction to nitric oxide by nitrite reductase, conversion of nitric oxide and ammonium to hydrazine by hydrazine synthase, and finally oxidation of hydrazine to dinitrogen gas by hydrazine dehydrogenase (Kartal et al., 2011b; Maalcke et al., 2016). The low potential electrons released upon the oxidation of hydrazine are postulated to be shuttled through a membrane embedded respiratory chain where they contribute to the *proton-motive force*, driving ATP synthesis (de Almeida et al., 2016). Finally, the electrons are shuttled back to the terminal electron acceptor nitrite.

Hydrazine synthase is a biochemically unique enzyme that produces hydrazine as a free metabolite in the anammox reaction (Kartal et al., 2011b). Crystallization of *K. stuttgartiensis* hydrazine synthase showed a crescent-shaped enzyme consisting of a dimer of αβγ-heterotrimers (Dietl et al., 2015). Each trimer holds one zinc ion, multiple calcium ions, and four iron-containing c-type hemes (αI, αII, γI, γII) of which two are postulated to have a catalytic function (αI and γI). Based on this structure, a two-step mechanism for hydrazine synthesis was proposed (Figure 1). In the first half-reaction of the proposed mechanism (Dietl et al., 2015), nitric oxide is reduced to hydroxylamine in the γ-subunit of hydrazine synthase. To perform this half-reaction, heme γII accepts electrons from an electron shuttle and transfers them to catalytic heme γI located 15 Å away. At catalytic heme γI, nitric oxide accepts three electrons and three protons, reducing nitric oxide to hydroxylamine. In the second half-reaction, hydroxylamine is proposed to combine with ammonia to hydrazine in the α-subunit of hydrazine synthase. For this reaction, hydroxylamine diffuses through an intra-enzymatic tunnel from the γ-subunit to catalytic heme αI. This transport is likely coordinated by a loop in the β-subunit of the enzyme. The function of the fourth heme, αII, is still unknown.

**Figure 1.**
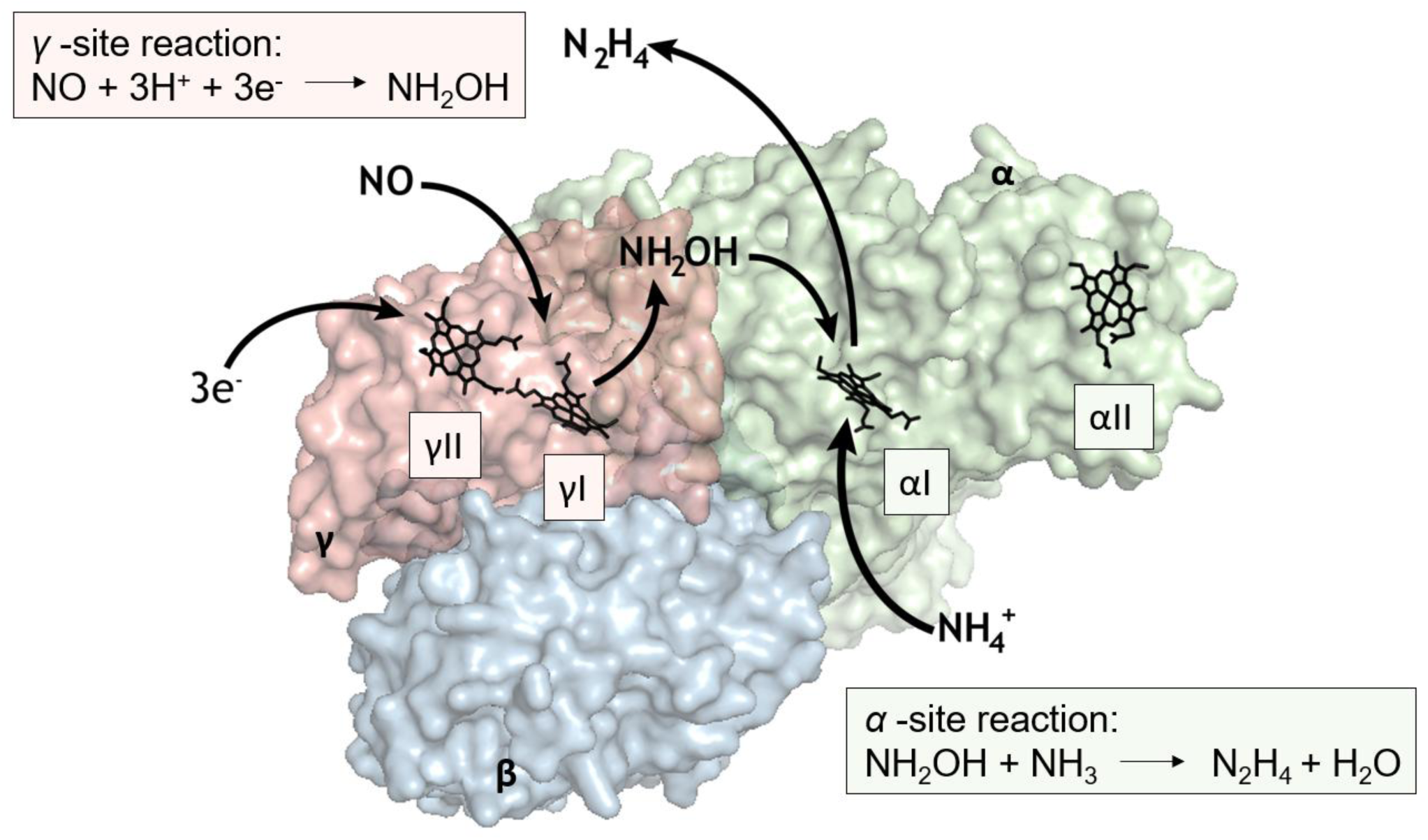
– Crystal structure of the hydrazine synthase αβγ-monomer isolated from anammox bacterium *Kuenenia stuttgartiensis* (PDB entry 5C2V) suggests a two-step reaction mechanism. Hydrazine synthase consists of a dimer of αβγ-heterotrimers. Each dimer contains four hemes proposed to aid in the formation of hydrazine from nitric oxide and ammonium. In the current working hypothesis, the surface exposed heme γII takes three electrons which are transferred to catalytic heme γI. There, nitric oxide is reduced to hydroxylamine (first half-reaction) which diffuses to the second catalytic heme: αI. At this heme, hydroxylamine and ammonia form hydrazine (second half-reaction). Figure inspired by Dietl et al. (2015).

Hydroxylamine is a potent inhibitor of hydrazine oxidation by the HAO-like protein hydrazine dehydrogenase ((Kartal et al., 2011b; Maalcke et al., 2016). Another HAO-like protein, hydroxylamine oxidoreductase (HOX), is therefore proposed to serve as a safety measure during the anammox reaction (Maalcke et al., 2014). HOX can oxidize hydroxylamine to nitric oxide. This way, HOX can scavenge and detoxify any potential hydroxylamine leaking from hydrazine synthase while at the same time providing nitric oxide as a substrate and three electrons required for the first half-reaction of hydrazine synthase. *In vitro*, HOX can be forced to also catalyze the oxidation of hydrazine to dinitrogen gas (Maalcke et al., 2014).

One way to experimentally verify the proposed two-step reaction mechanism of hydrazine synthase is via enzymatic assays. However, hydrazine cannot be measured directly due to its reactive nature and low molecular weight (32.0452 g/mol). Previously, enzymatic hydrazine production was measured indirectly using a coupled enzyme assay with isolated hydrazine synthase incubated with nitric oxide and ammonium. HOX, capable of oxidizing hydrazine at 1.6 µmol/min/mg protein (Maalcke et al., 2014), was used to convert hydrazine to dinitrogen gas. In this assay, bovine cytochrome *c* serves both as an electron donor to reduce hydrazine synthase heme γII and as an electron acceptor to oxidize HOX. Assuming hydrazine synthesis is the rate-limiting step, a rate of 0.3 nmol dinitrogen gas/min/mg protein can be deduced as the hydrazine synthesis rate (Kartal et al., 2011b). Whole *K. stuttgartiensis* cells can produce up to 30 nmol dinitrogen gas/min/mg protein (Kartal et al., 2011b), indicating that cellular activity is 100 times higher than the activity of isolated hydrazine synthase.

The loss of most hydrazine synthesis activity during cell disruption might indicate a limiting factor in the assay with isolated hydrazine synthase. As hydrazine synthase was aerobically isolated in these previous experiments and its native environment is anaerobic, oxygen may have damaged the enzyme resulting in reduced activity (Kartal et al., 2011b). Anaerobically isolated hydrazine synthase (Versantvoort et al., 2025) showed noticeable differences in its EPR spectrum compared to the aerobically isolated enzyme (Dietl et al., 2015), indicating oxygen may damage the enzyme resulting in reduced activity. Alternatively, isolated hydrazine synthase could also be slow due to the disruption of a multi-component enzyme system in which hydrazine production is the rate-limiting step (Kartal et al., 2011b), or the absence of a suitable electron donor. In addition to these limitations in enzyme activity, the coupled assay is unsuitable to measure the proposed second half-reaction of hydrazine production because HOX uses hydroxylamine as substrate (Maalcke et al., 2014). Therefore, optimization of both the hydrazine synthase isolation procedure and enzyme assay is key to further our understanding of hydrazine synthase functioning.

Here, we investigated the molecular mechanism and activity of *K. stuttgartiensis* MBR1 hydrazine synthase using an optimized anaerobic isolation method, to minimize activity loss, and a direct LC-MS hydrazine quantification method instead of the coupled enzyme assay. We determined kinetic parameters and set out to verify the hypothesized second half-reaction in which hydroxylamine and ammonium yield hydrazine.

## Materials & methods

### *Cultivation of* K. stuttgartiensis *cells*

*“Candidatus* Kuenenia stuttgartiensis*”* strain MBR1 (Frank et al., 2018) was cultured as planktonic cells in a 12 L membrane bioreactor (MBR) at an OD_600_ of ∼1.2 (Applikon B.V. Schiedam, The Netherlands), as described by Kartal et al. (2011a). In brief, the bioreactor was kept anoxic by continuous flushing of the bioreactor itself and medium vessel with argon/carbon dioxide (95/5%, 10 mL/min). Carbon dioxide in the supplied gas was sufficient to maintain the pH in the bioreactor between 7.0 and 7.4. Planktonic biomass was continuously removed at 1.1 L per day, under constant sparging with dinitrogen gas to maintain anaerobicity, for preparation of cell-free extract. The medium consisted of: 45 mM (NH_4_)_2_SO_4_, 45 mM NaNO_2_, 10 mM KHCO_3_, 0.2 mM NaH_2_PO_4_, 0.6 mM HCl, 1 mM CaCl_2_·2H_2_O, 0.4 mM MgSO_4_·7H_2_O and 22.5 µM FeSO_4_·7 H_2_O, and trace elements 0.625 µM CoCl_2_·6H_2_O, 0.625 µM CuSO_4_·5H_2_O, 0.125 µM H_3_BO_3,_ 3.125 µM MnCl_2_·4H_2_O, 0.563 µM Na_2_MoO_4_·2H_2_O, 0.125 µM Na_2_WO_4_·2H_2_O, 0.5 µM NiCl_2_·6H_2_O, 0.375 µM SeO_2_, 25.188 µM C_10_H_14_N_2_Na_2_O_8_·2H_2_O, and 0.938 nM ZnSO_4_·7H_2_O.

### Preparation of the soluble protein fraction, crude hydrazine synthase fraction and isolated hydrazine synthase fraction from K. stuttgartiensis cells

Procedures were optimized from Kartal et al. (2011b), Maalcke et al. (2014), and Akram et al. (2019) with the most important difference that all transfer and isolation steps were carried out at 16°C under anaerobic conditions in a glovebox atmosphere (or closed vessel for transport to French press) containing 100% dinitrogen gas (Versantvoort et al., 2025). In brief, 700 mL collected culture was transferred to centrifugation tubes with a rubber seal. *K. stuttgartiensis* cells were pelleted by centrifugation at 10.000 x *g* for 10 minutes at 4°C (Sorvall centrifuge, fixed angle). The cell pellet was resuspended in 10 mL 20 mM Tris, pH 7.3. Cells were broken by passing them once through a French press cell at 138 MPa (American Instrument Company). The broken cells were collected in an anaerobic bottle with a rubber seal and transferred to ultra-centrifuge tubes with a rubber seal. Membranes and whole cells were pelleted by centrifugation at 162.000 x *g* for one hour at 4°C (Optima-XE-90, Fixed-Angle 90 Ti Rotor, Beckman Coulter) resulting in the soluble protein fraction in the supernatant. The soluble protein fraction was either buffer exchanged to 20 mM MOPS, 100 mM NaCl, pH 7.5 and concentrated using 30 kDa spinfilters (Sartorius) to be used directly in experiments or further separated to obtain crude hydrazine synthase fraction and isolated hydrazine synthase.

To obtain crude hydrazine synthase fraction, 10 mL soluble protein fraction was loaded onto a Q-Sepharose column (XK 26/20, GE Healthcare/Cytiva, 70 mL, packed in house). This crude hydrazine synthase fraction contained HOX (KSMBR1_2670/kustc1061, Genbank CAJ71806) (Maalcke et al., 2014), the electron/NO shuttle NaxLS (KSMBR1_3082-83/kusta0087-88, Genbank SOH05560-61) (Akram et al., 2019), nitrite oxidoreductase (NXR, KSMBR1_1455 and KSMBR1_1458-59/kustd1700 and kustd1703-04, Genbank SOH03954, SOH03957-58) (Chicano et al., 2021) and hydrazine synthase (KSMBR1_2711-13 and KSMBR1_3601-03/kuste2859-61, GenBank CAJ73611-13) (Versantvoort et al., 2025). The column was connected to an ÄKTA purifier (GE Healthcare/Cytiva), equilibrated with 20 mM Tris, pH 7.3 and run with a flowrate of 5 mL/min. The crude hydrazine synthase fraction was eluted isocratically with 20 mM Tris, 200 mM NaCl, pH 7.3. The fraction was either buffer exchanged to 20 mM MOPS buffer, pH 7.0 and concentrated using 30 kDa spinfilters (Sartorius) to be used directly in experiments or used to isolate hydrazine synthase, HOX, NaxLS, and NXR.

To separate the proteins in the crude hydrazine synthase fraction, the fraction was buffer-exchanged to 20 mM potassium phosphate buffer, pH 7.0 and concentrated to 10 mL using a 50 kDa cutoff Amicon pressure filter unit. The sample was loaded onto a Hydroxyapatite column (CHT type II, 40 µm particle size, GE Healthcare/Cytiva, 15 mL, packed in house), equilibrated with 20 mM potassium phosphate buffer, pH 7.0, run with a flowrate of 5 mL/min. Proteins were eluted in a linear gradient from 20-500 mM potassium phosphate. NaxLS eluted first in the flow-through, then HOX eluted at 50 mM (Maalcke et al., 2014), hydrazine synthase at 150 mM (Kartal et al., 2011b), and NXR at 200 mM potassium phosphate. The protein fractions were concentrated using 30 kDa spinfilters (Sartorius).

Hydrazine synthase concentration in µg/µL was calculated from the absorbance at a wavelength of 280 nm measured with a Cary 60 spectrophotometer (Agilent) and calculated with the formula: concentration (µg/µL) = (A_280_ / ε_280_ of hydrazine synthase) × molecular weight of hydrazine synthase. In which the 280 nm extinction coefficient is 26,7250 M^-1^cm^-1^, and the molecular weight of hydrazine synthase is 160,997.73 Da. For the other proteins, the concentration in µg/µL was calculated from the absorbance measured at wavelengths 260 nm and 280 nm. Protein concentrations were calculated with the formula: concentration (µg/µL) = A_280_ × 1.55 – A_260_ × 0.76 (Stepanchenko et al., 2011). All protein samples were aliquoted and stored anaerobically at –20°C until used.

Purity of isolated hydrazine synthase samples was verified via typical absorption spectra of sodium dithionite-reduced hydrazine synthase as previously described (Dietl et al., 2015; Versantvoort et al., 2025). For a more accurate assessment, sodium dodecyl sulfate polyacrylamide gel electrophoresis (SDS-PAGE) (10% resolving and 4% stacking gel) was used (Supplementary figure 1) (Laemmli, 1970). To confirm the absence of HOX in the hydrazine synthase sample, absorption at 460 nm in a sodium dithionite-reduced sample was assessed spectrophotometrically (Maalcke et al., 2014). Additionally, the concentration of HOX was estimated from the hydrazine oxidation rate. Pure HOX oxidizes hydrazine with a specific activity of 1.6 µmol/min/mg (Maalcke et al., 2014). The highest rate observed in our samples was 2.4 nmol/min/mg, indicating a maximum of 0.15% HOX.

### Activity assays: hydrazine production from ammonium and nitric oxide measured via coupled assays

#### GC-MS coupled assay with the soluble protein fraction

To evaluate the effect of oxygen on hydrazine production by the soluble protein fraction, gas chromatography mass spectrometry (GC-MS) assays with anaerobically prepared the soluble protein fraction exposed to ambient air were executed. In this activity assay, reduced hydrazine synthase produces hydrazine from nitric oxide and ammonium, and HAO-like proteins convert hydrazine to dinitrogen gas. The dinitrogen gas formation was followed over time via GC-MS (Kartal et al., 2011b).

GC-MS assays were performed in 7-mL UV-protected Amber glass vials with butyl stoppers and a metal cap. All additions and gas sampling were done with gas tight syringes (Hamilton). All reagents and stock solutions were made anoxic in three cycles of 10 minutes in which vacuum and sparging with argon were alternated. Reaction mixtures consisted of 50 µM sodium ascorbate (L-ascorbic acid, Sigma life sciences) as electron donor and 1 mM ^15^N-labeled ammonium chloride (99%, Cambridge Isotope Laboratories) in an end volume of 1 mL 20 mM MOPS, 100 mM NaCl buffer, pH 7.5 with a headspace of 0.4% nitric oxide. The reaction was started with the addition of 5 mg soluble protein fraction and 1 mM phenazine methosulphate as electron carrier (Acros organics). Oxygen-exposed soluble protein fraction was obtained by exposing anaerobically isolated soluble protein fraction to ambient air for 15 minutes prior to the assay. After air exposure, the sample was made anaerobic again as specified above. To measure hydrazine production, ^15^N^14^N-labeled dinitrogen gas formation in the headspace was followed over time with an Agilent 6890 series gas chromatograph coupled to a 5975C inert MSD mass spectrometer equipped with a Porapak Q column heated to 80°C with helium as the carrier gas. Calibration curves were generated by triplicate injections of different volumes of a known ^15^N^15^N-dinitrogen gas concentration.

#### MIMS coupled assay with crude hydrazine synthase fraction or isolated hydrazine synthase

To test reproducibility of hydrazine synthesis and the specific hydrazine synthase activity in crude hydrazine synthase fraction, coupled activity assays were executed using membrane inlet mass spectrometry (MIMS). In these assays, reduced hydrazine synthase produced hydrazine from added ammonium and nitric oxide and HOX converted hydrazine to dinitrogen gas. The dinitrogen gas formation was followed continuously over time with MIMS (HPR40, Positive Ion Counting detector, Hiden Analytical). The coupled assays were performed in a custom-built chamber (Schmitz et al., 2020) with an 8 mm^2^ MIMS probe covered with a 10 µm thick silicon membrane. The chamber was filled with buffer and made anaerobic by flushing it with Argon for 30-60 minutes. The chamber was closed resulting in a reaction volume of 4-5 mL. All additions of reagents and proteins from anoxic stock solutions were made through a capillary using gastight syringes.

For the measurement of hydrazine synthesis reproducibility in crude hydrazine synthase fraction (eluted with 200 mM NaCl from the Q-Sepharose column), the reaction mixture consisted of 10.9 mg crude hydrazine synthase fraction, 50 µM bovine cytochrome c (Sigma Aldrich) as electron donor and acceptor, and 235 µM ^15^N-ammonium. The reaction was started with the addition of 28 µM hydrazine (99%, Sigma Aldrich). Sequential additions of nitric oxide were made from ultrapure water (PURELAB Chorus 1, Veolia) bubbled with 4% nitric oxide. The assay was carried out in 20 mM MOPS, pH 7.0.

To measure specific hydrazine synthase activity in the crude hydrazine synthase fraction, the four proteins of the crude fraction (hydrazine synthase, NaxLS, NXR and HOX) were added to the reaction mixture in known concentrations after their additional isolation on a Hydroxyapatite column. The reaction mixture consisted of 450 µg hydrazine synthase, 100 µg NaxLS, and 100 µg NXR, 50 µM bovine cytochrome c as electron donor and acceptor, 400 µM ^15^N-ammonium, and 46 µM nitric oxide. This higher concentration of nitric oxide was chemically formed from 40 µM sodium dithionite (Merck) and 40 µM sodium nitrite (Sigma Aldrich). The reaction was started with 270 µg HOX and carried out in 50 mM HEPES, pH 7.5. 25 min after HOX addition, an additional oxidized 50 µM cytochrome c was injected. To determine the specific activity of hydrazine synthase, the rate of ^15^N^14^N-dinitrogen gas production was divided by the amount of hydrazine synthase in the assay.

### Activity assays: direct LC-MS assays with hydrazine synthase incubated with ammonium and hydroxylamine

To measure the proposed second half-reaction of hydrazine synthesis, hydrazine formation from ammonium and hydroxylamine by isolated hydrazine synthase was followed over time with liquid chromatography mass spectrometry (LC-MS). Typically, activity assays were carried out under anaerobic conditions at 16°C in an end volume of 500 µL 20 mM potassium phosphate buffer, pH 7.0 in open UV-protected glass vials.

To measure hydrazine synthase activity, the reaction mixture contained 150 µg hydrazine synthase with 1 mM ammonium chloride (Fisher Chemical) and 1 mM hydroxylamine hydrochloride (99%, Thermo Scientific). Controls contained either 150 µg inactivated hydrazine synthase with 1 mM ammonium and 1 mM hydroxylamine hydrochloride, only 150 µg hydrazine synthase, only 1 mM ammonium and 1 mM hydroxylamine, or 150 µg hydrazine synthase with either 1 mM ammonium or 1 mM hydroxylamine. Hydrazine synthase was inactivated by heating the protein sample to 95°C for 20 minutes resulting in a light red and turbid sample. Additionally, hydrazine synthesis was measured in reaction mixtures containing 15 or 60 µg hydrazine synthase and 10 µM hydroxylamine and 1 mM ammonium.

To evaluate assay reproducibility, effects of pH on hydrazine synthesis, potential product inhibition, and the effect of oxygen on hydrazine synthesis were examined. The reaction mixture contained 60 µg hydrazine synthase, 10 µM hydroxylamine and 1 mM ammonium. Assays to evaluate the effect of pH were carried out in 20 mM potassium phosphate buffer, pH 6.0, 7.0, and 8.0. Potential product inhibition was tested by addition of 0.5-5 µM ^15^N^15^N-labeled hydrazine (99%, Cambridge Isotope Laboratories) to the reaction mixture prior to activity measurements. To measure the effects of oxygen, anaerobically isolated hydrazine synthase was exposed to ambient air for 15 minutes prior to activity measurements or the activity assay was carried out in aerobic conditions (ambient air) at 20°C. Enzyme kinetic parameters were evaluated with 60 µg hydrazine synthase incubated with various concentrations of hydroxylamine and ammonium ranging between 0 and 10 mM.

To measure whether hydrazine synthase was inactivated during the assay, the reaction was initiated with 60 µg hydrazine synthase, 10 µM hydroxylamine and 1 mM ammonium to which an additional 30 µg hydrazine synthase was added six minutes after the assay started.

Hydrazine synthesis was measured as described in (Vermeir et al., 2026). In brief, 25 µL reaction mixture and 25 µL internal standard (^15^N^15^N-labeled hydrazine) were added to 50 µL derivatizing solution. Samples were incubated in the dark for at least one hour for the derivatization of hydrazine with benzaldehyde to form 1,2-dibenzylidenehydrazine and were either measured the same day or stored at –70°C until measurement. Prior to transferring samples to autosampler vials for LC-MS measurements, they were centrifuged at 20817 x *g* for five minutes at 15°C to remove denatured hydrazine synthase.

Control samples were prepared during every assay. To this end, 1 µL of hydrazine synthase stock solution, 99 µL 20 mM potassium phosphate buffer, pH 7.0 supplemented with hydroxylamine and ammonium, and 100 µL internal standard were added to 200 µL derivatizing solution. For activity assays without hydrazine synthase, 20 mM potassium phosphate buffer, pH 7.0 was added instead.

Samples were subjected to LC-MS analysis performed by an Agilent 1290 II LC system with a 1290-series isocratic pump, multisampler, MCT, and high-speed pump coupled to an Agilent Accurate Mass 6546 Quadrupole Time of Flight (Q-TOF) instrument operated in the positive ionization mode. Sample vials with PTFE/silicone septa (Agilent) were used and the autosampler was set at 8°C. The isocratic pump continuously pumped a reference solution consisting of 1 µM Hexakis (1H, 1H, 3H-tetrafluoropropoxy) phosphazine, 5 µM purine, and 25 µM ammonium trifluoroacetate (Agilent technologies) at 2.5 mL/min, of which 1/100 was directed to the source via a splitter. The MS signals produced by reference compounds Hexakis and purine (m/z 121.0508 and 922.0997, respectively) were used for continuous mass calibration.

In initial experiments, derivatized hydrazine was measured using aqueous normal phase chromatography as described by Jansen et al. (2020). In brief, a Cogent Diamond Hydride column (2.1 × 150 mm, 4 μm, Cogent HPLC Columns) equipped with a 0.3 µm inline filter and 0.3 μm frit (Agilent Technologies) was used at 25°C and with a maximum pressure of 400 bar. A sample volume of 2 µL was injected onto the column followed by separation with a 0.4 mL/min gradient of water (A) and acetonitrile (B) (both with 0.2% formic acid (Honeywell Fluka): 85% B from 0-2 minutes, 80% B from 3-5 minutes, 75% B from 6-7 minutes, 70% B from 8-9 minutes, 50% B from 10-11 minutes, 20% B from 11.10-14 minutes, 5% B from 14.10-24 minutes. Followed by re-equilibration at 85% B for 10 minutes. The LC stream was directed to the MS from 0.5 to 3.5 minutes and to waste during the rest of the runs.

For reversed phase chromatography, samples were subjected to LC-MS analysis via the method described in (Vermeir et al., 2026). In brief, 3 µL sample was injected onto a Poroshell 120 EC-C18 column (2.1 × 50 mm, 1.9 μm, Agilent Infinity Lab), followed by separation with a 0.4 mL/minute gradient of water (A) and acetonitrile (B) (both with 0.2% formic acid (Honeywell Fluka)).

MS spectra were collected using full scan with a range of m/z 50-1200 at 1000 ms/spectrum. Collected data were analyzed using Quant software 10.0 from Agilent (Agilent Technologies). Signals for hydrazine and ^15^N^15^N-hydrazine (m/z 209.1074 and m/z 211.1014, respectively), were analyzed with a 100-ppm window and peak smoothing (function width 15, Gaussian width 5).

For calibration curves, 25 µL hydrazine stock solution and 25 µL internal standard (^15^N^15^N-hydrazine) were mixed with 50 µL derivatizing solution. The calibrator levels were vortexed for 10 seconds and incubated for one hour in the dark at room temperature. Blanks consisted of derivatizing solution diluted with ultrapure water (1:1 (v/v)). For the quantification of hydrazine in samples to which ^15^N^15^N-hydrazine was added to the reaction mixture of the assay, ^15^N^15^N-hydrazine was not considered as internal standard for the calibration curve or for the samples measured.

### Calculations and figures

ANOVA one-way tests were performed in Excel. Hydrazine synthesis rates were calculated from MIMS data using linear regression on the initial linear part of the curve, analyzed with OriginPro 2020b (OriginLab). To determine hydrazine synthesis rates in direct assays, the initial activity burst measured in activity assays was excluded and only the linear part of the curve was used. Figures were made in Pymol 3.0 (Schrödinger et al., 2020) from the hydrazine synthase crystal structure PDB entry 5C2V (Dietl et al., 2015) and in Rstudio version 4.4.1 with the readxl, ggplot2, ggsignif, drc, and cowplot packages.

## Results

### Hydrazine synthesis by K. stuttgartiensis soluble protein fraction is negatively affected by oxygen exposure prior to the coupled activity assay

To evaluate the impact of oxygen on hydrazine synthase activity, we measured activity in an anaerobically prepared soluble protein fraction kept under anaerobic conditions, as well as after exposing the fraction to ambient air prior to performing a coupled assay. Both coupled assay activity measurements were carried out under anaerobic conditions. Hydrazine synthesis was followed indirectly via oxidation of hydrazine to dinitrogen gas by HAO-like proteins using GC-MS (Figure 2). Hydrazine synthesis is assumed to be the rate-limiting step in this process. After exposure to ambient air, the soluble protein fraction produced significantly less dinitrogen gas (Table 1). Since HAO-like proteins remain active when isolated under aerobic conditions (Maalcke et al., 2014; Maalcke et al., 2016), hydrazine oxidation to dinitrogen gas is most likely unaffected. This suggests that oxygen can impact hydrazine synthase activity, which prompted us to isolate hydrazine synthase under strict anaerobic conditions for further experiments.

**Figure 2.**
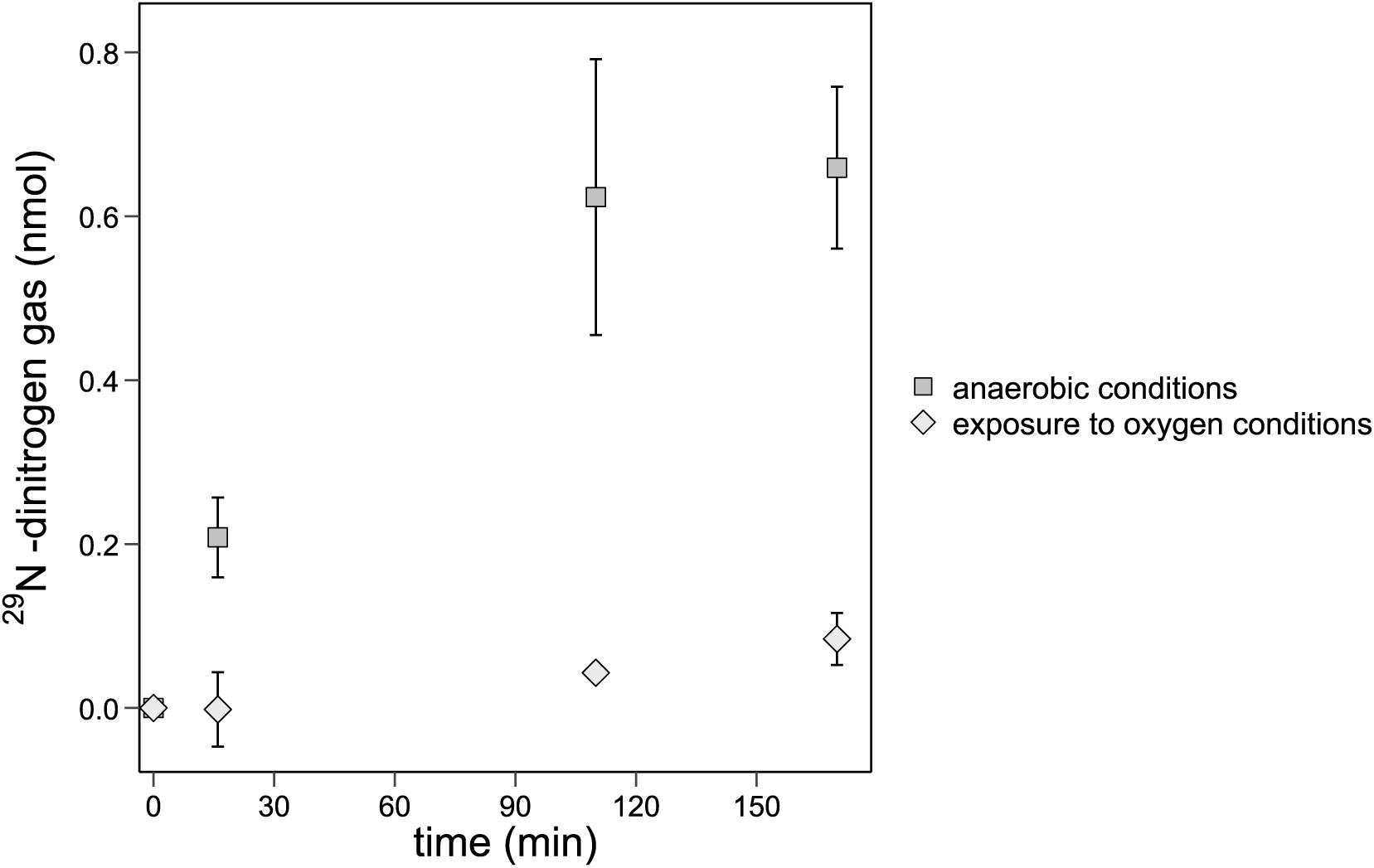
– Hydrazine synthesis in anaerobically prepared soluble protein fraction from *K. stuttgartiensis* measured after proteins were exposed to ambient air for 15 minutes and in proteins kept under strictly anaerobic conditions. Ambient air exposure reduces hydrazine synthesis by the soluble protein fraction. Activity assays contained 50 µM sodium ascorbate as electron donor, 1 mM phenazine methosulphate as electron carrier, 1 mM ^15^N-labeled ammonium, and 5 mg soluble protein fraction in 1 mL 20 mM MOPS 100 mM NaCl buffer pH 7.5 with a 0.4% nitric oxide headspace. Assays were carried out in anaerobic conditions. Enzyme activity was measured with GC-MS via the enzymatic conversion of the produced hydrazine to dinitrogen gas. Data represented as the mean ± SD (*n*=2 technical replicates).

**Table 1.** – Overview of hydrazine synthase (HZS) activity in different experiments measured via the coupled enzyme assay. * Data from Kartal et al. (2011b). ** Aerobic = after exposure to ambient air for 15 min.

| Conditions in which HZS activity was measured | Average HZS activity ± SD |  | Average HZS activity ± SD |  |
| --- | --- | --- | --- | --- |
|  | (nmol hydrazine/min/mg protein) | Figure | (nmol <sup>15</sup>N<sup>14</sup>N-dinitogen gas) | Figure |
| 5 mg soluble protein fraction + 1 mM <sup>15</sup>N-labelled ammonium<br>+ 0.4% nitric oxide headspace + 50 µM sodium ascorbate<br>+ 1 mM phenazine methosulphate |  |  |  | 2 |
| anaerobic soluble protein fraction |  |  | 0.7 ± 0.10 |  |
| aerobic** soluble protein fraction |  |  | 0.1 ± 0.03 |  |
| 10.9 mg crude HZS fraction + 235 µM <sup>15</sup>N-labelled ammonium<br>+ 50 µM bovine cytochrome c |  | 3 |  |  |
| extra addition of 0.56 µM nitric oxide | 0.03 ± 0.001 |  |  |  |
| extra addition of 0.56 µM nitric oxide | 0.04 ± 0.001 |  |  |  |
| extra addition of 0.28 µM nitric oxide | 0.04 ± 0.004 |  |  |  |
| extra addition of 0.84 µM nitric oxide | 0.05 ± 0.001 |  |  |  |
| 270 µg HOX + 450 µg HZS + 100 µg NaxLS + 100 µg NXR<br>+ 40 µM sodium dithionite + 10 µM sodium nitrite<br>+ 50 µM bovine cytochrome c + 400 µM <sup>15</sup>N-labelled ammonium<br>+ 6 µM nitric oxide | 0.03 | 4 |  |  |
| extra addition of 50 µM bovine cytochrome c | 22.00 |  |  |  |
| whole *K. stuttgartiensis* cells + nitrite + ammonium | 30.0 | * |  |  |
| cell free extract from *K.stuttgartiensis* |  |  |  |  |
| nitrite + ammonium | 0 | * |  |  |
| nitric oxide + ammonium | 0.06 | * |  |  |
| 1.6 mg HZS + 4.7 ug HOX + 1 mM <sup>15</sup>N-labelled ammonium +<br>0.9 mM nitric oxide<br>+ 50 µM bovine cytochrome c | 0.3 | * |  |  |

### The hydrazine synthesis rate in crude hydrazine synthase fraction was reproducible but its measurement via the coupled assay was influenced by electron donor and acceptor availability

Next, we tested the reproducibility of hydrazine synthesis in an anaerobically isolated crude hydrazine synthase fraction. This fraction eluted with 200 mM NaCl from the Q-Sepharose column and is primarily composed of hydrazine synthase, HOX, the electron/nitric oxide shuttle NaxLS, and nitrite oxidoreductase (NXR). In this assay, ^15^N^14^N-labelled hydrazine synthesized from ^14^N-nitric oxide and ^15^N-labeled ammonium by hydrazine synthase would be rapidly oxidized to ^15^N^14^N-labeled dinitrogen gas by HOX. Hydrazine synthesis is considered the rate-limiting step, and the ^15^N^14^N-dinitrogen gas formation was followed continuously with MIMS as a proxy for hydrazine production (Figure 3). When the crude hydrazine synthase fraction was incubated with ^15^N-ammonium and ^14^N^14^N-hydrazine in the absence of nitric oxide, some ^15^N^14^N-dinitrogen gas was formed. Possibly, residual nitric oxide was bound to hydrazine synthase during enzyme isolation (Versantvoort et al., 2025) and served as a substrate for enzymatic hydrazine synthesis. Upon additions of ^14^N-nitric oxide, ^15^N^14^N-dinitrogen gas was produced. As expected, the amount of ^15^N^14^N-dinitrogen gas production was dependent on the amount of nitric oxide added. Moreover, all nitric oxide seemed to be converted (Figure 3). Despite varying nitric oxide concentrations added, the rate remained similar: ∼0.04 nmol ^15^N^14^N-dinitrogen gas/min/mg protein, indicating that the *K_m_* value of hydrazine synthase is below 0.3 µM nitric oxide (Table 1). Interestingly, besides production of ^15^N^14^N-dinitrogen gas, a small amount of ^15^N^15^N-dinitrogen gas was produced, amounting to 10% of the ^15^N^14^N-dinitrogen gas production, with ^15^N from ammonium being the only heavy nitrogen isotope present.

**Figure 3.**
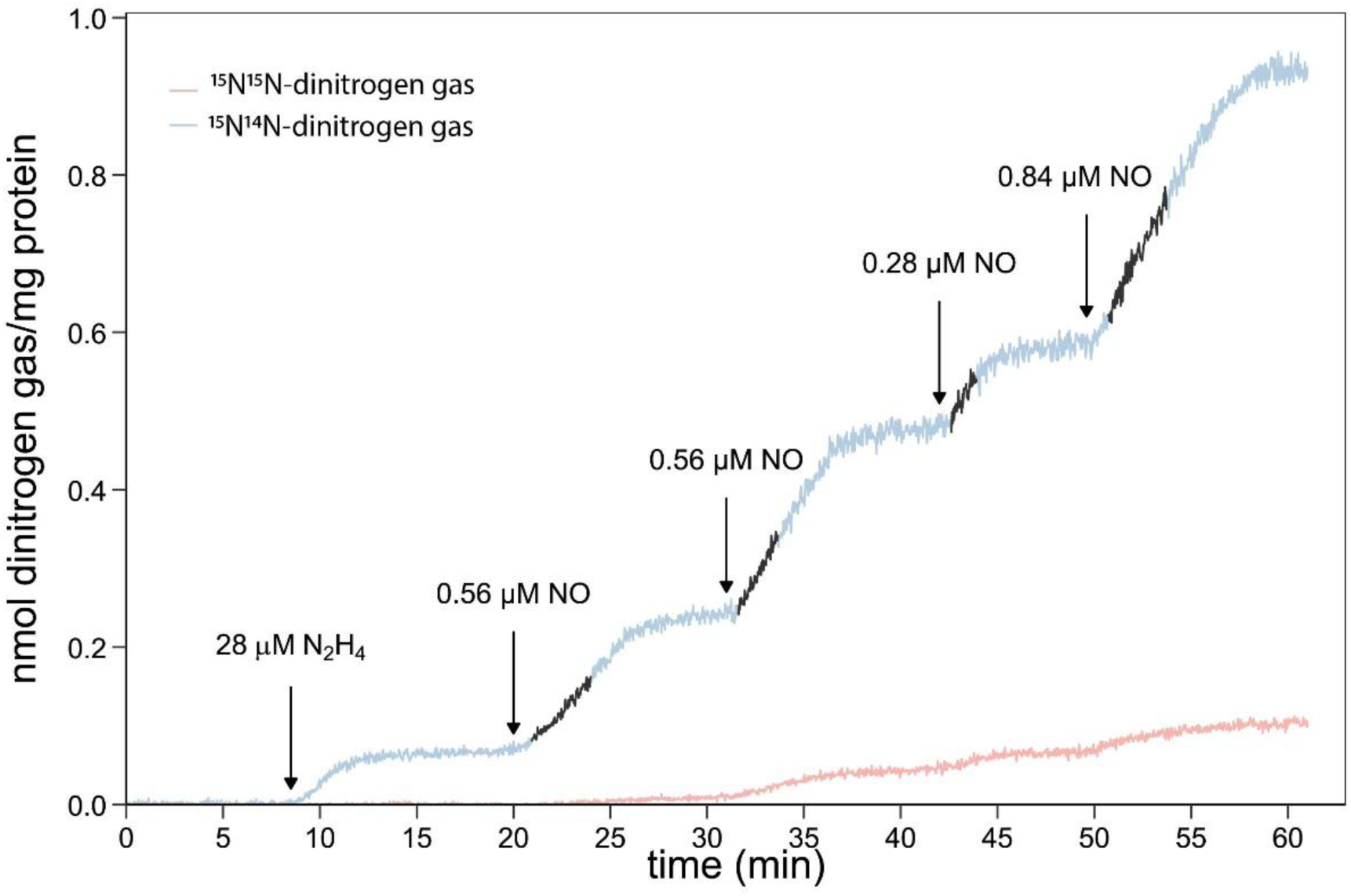
– Hydrazine synthesis in *K. stuttgartiensis* crude hydrazine synthase fraction measured via ^15^N^14^N-labeled dinitrogen gas with various concentrations of nitric oxide. The results indicate that all nitric oxide is converted to hydrazine, hydrazine synthesis is reproducible, and the rate is independent of nitric oxide concentrations above 0.28 µM. The activity assay contained 10.9 mg crude hydrazine synthase fraction, 235 µM ^15^N-ammonium with 50 µM bovine cytochrome c as electron donor and acceptor in 20 mM MOPS buffer, pH 7.0. The reaction was started with the addition of 28 µM hydrazine. Enzyme activity was measured with MIMS via the enzymatic conversion of hydrazine to dinitrogen gas. Data points included to calculate the hydrazine synthesis rate are indicated in black (*n*=1).

To investigate the specific rate of hydrazine synthesis in the crude hydrazine synthase fraction, the four proteins of the crude fraction were added in known concentrations after additional isolation. Initial specific hydrazine synthesis was 0.03 nmol/min/mg hydrazine synthase (Figure 4, Table 1), similar to the calculated rate for the crude hydrazine synthase fraction (see above). In an attempt to increase the rate, oxidized cytochrome c was added after 35 minutes. This resulted in a boost of ^15^N^14^N-dinitrogen gas production and a calculated hydrazine synthesis rate of 22 nmol/min/mg hydrazine synthase (Figure 4, table 1). However, it is uncertain whether hydrazine is produced at this rate as hydrazine is not measured directly in this coupled assay, potentially resulting in a pool of hydrazine accumulating due to a shortage of oxidized cytochrome c, preventing HOX from fully converting hydrazine to dinitrogen gas. Consequently, the production of dinitrogen gas becomes limited by the available oxidized cytochrome c, potentially skewing the measured hydrazine activity rates. Over 40 minutes, 65 nmol ^15^N^14^N-dinitrogen gas/mg protein was produced, indicating a minimum rate for hydrazine synthesis of 1.5 nmol/min/mg protein. In conclusion, accurately determining hydrazine synthase rates via the coupled enzyme assay requires precisely balancing the electron donor pool for hydrazine synthesis and the electron acceptor pool for hydrazine oxidation to avoid over– or underestimation. This is, however, technically challenging.

**Figure 4.**
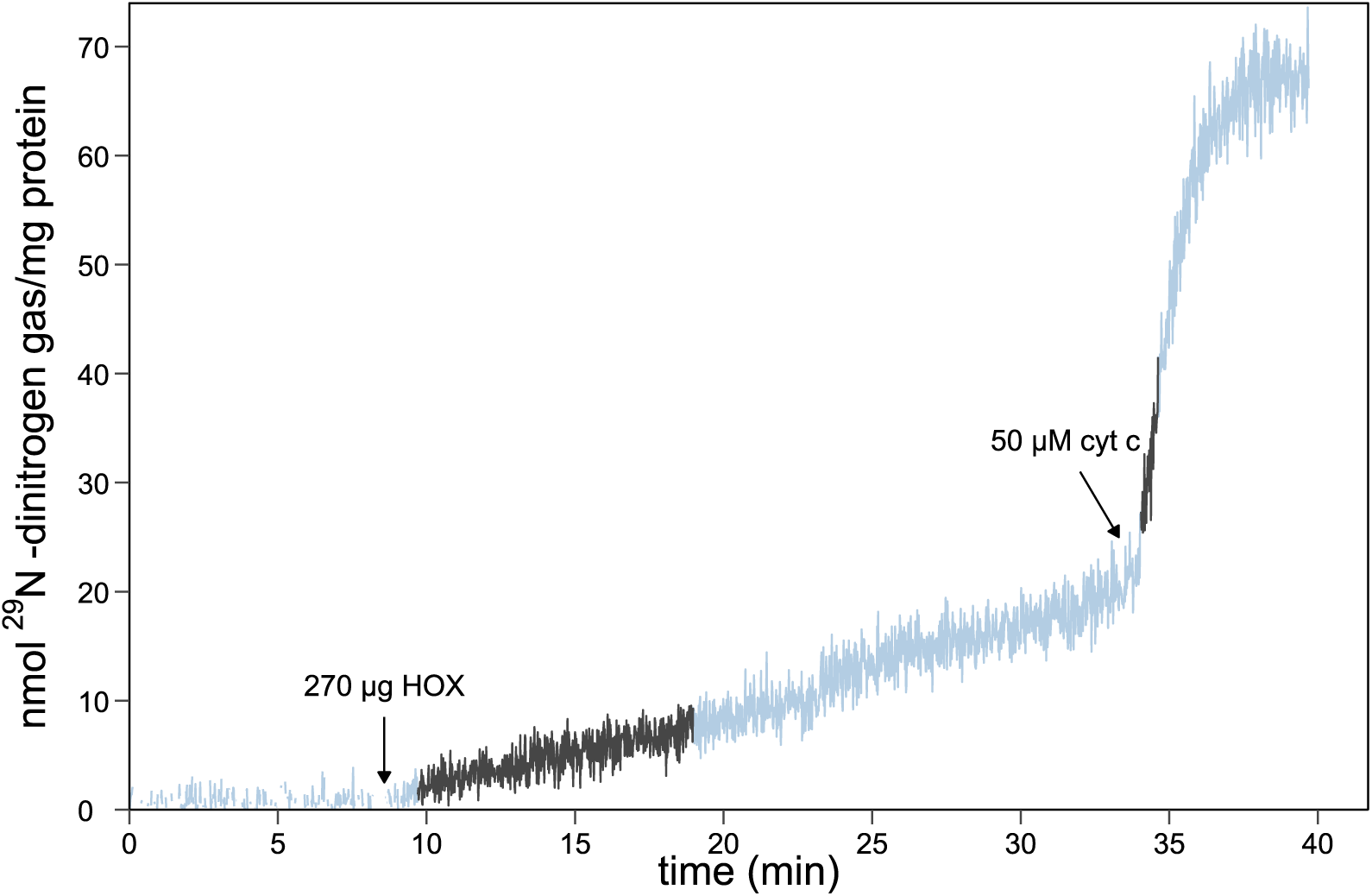
– Hydrazine synthesis measured via the coupled enzyme assay with HOX to which extra cytochrome c was added during the assay. Oxidized cytochrome c availability influences hydrazine synthesis measurements in the coupled assay with hydrazine synthase and HOX. The activity assay contained 450 µg hydrazine synthase, 50 µM bovine cytochrome c as electron donor and acceptor, 400 µM ^15^N-ammonium, 6 µM nitric oxide, 100 µg NaxLS, and 100 µg NXR. The reaction was started with HOX and carried out in 50 mM HEPES pH 7.5. Enzyme activity was measured with MIMS via the enzymatic conversion of hydrazine to dinitrogen gas. Data points included to calculate the hydrazine synthesis rate are indicated in black (*n*=1).

### Hydrazine synthase can synthesize hydrazine from hydroxylamine and ammonium

To circumvent the challenges in electron balancing encountered in the coupled enzyme assay, we developed a method in which hydrazine produced by isolated hydrazine synthase is chemically derivatized and subsequently measured with LC-MS (Vermeir et al., 2026). With this method, we investigated whether hydrazine synthase performs the proposed second half-reaction in which hydroxylamine and ammonium yield hydrazine. Hydrazine synthase produced 0.46 ± 0.06 nmol hydrazine/min/mg protein from ammonium and hydroxylamine (Figure 5, Table 2). Control experiments without hydrazine synthase, without substrates or with inactivated hydrazine synthase confirmed that hydrazine formation was enzymatic. Hydrazine synthase incubated solely with hydroxylamine exhibited minimal activity with a rate of 0.03 nmol hydrazine/min/mg protein. Thus, isolated hydrazine synthase can use externally supplied hydroxylamine and ammonium to form hydrazine.

**Figure 5.**
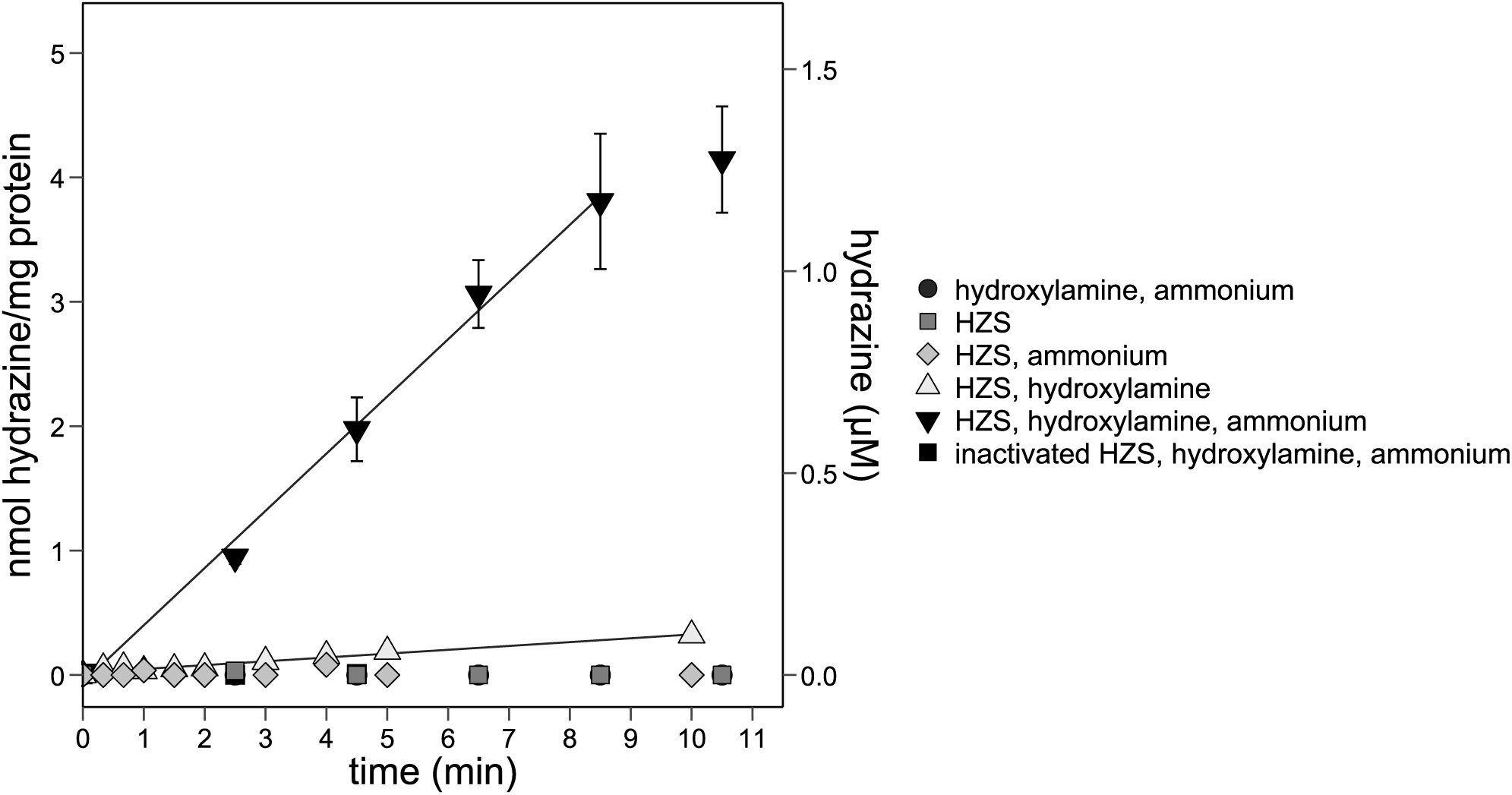
– Hydrazine synthase can synthesize hydrazine from hydroxylamine and ammonium. Hydrazine synthase produced hydrazine from 1 mM externally supplied hydroxylamine and 1 mM ammonium. Control experiments without hydrazine synthase, without substrates or with inactivated hydrazine synthase confirmed that hydrazine formation was enzymatic. When hydrazine synthase was incubated with hydroxylamine alone, some hydrazine was produced. Data points included to calculate the hydrazine synthesis rate are connected by a trendline. Data are presented as mean ± SD (*n=3* technical replicates, except for assays with only ammonium or only hydroxylamine: *n=1*).

**Table 2.**
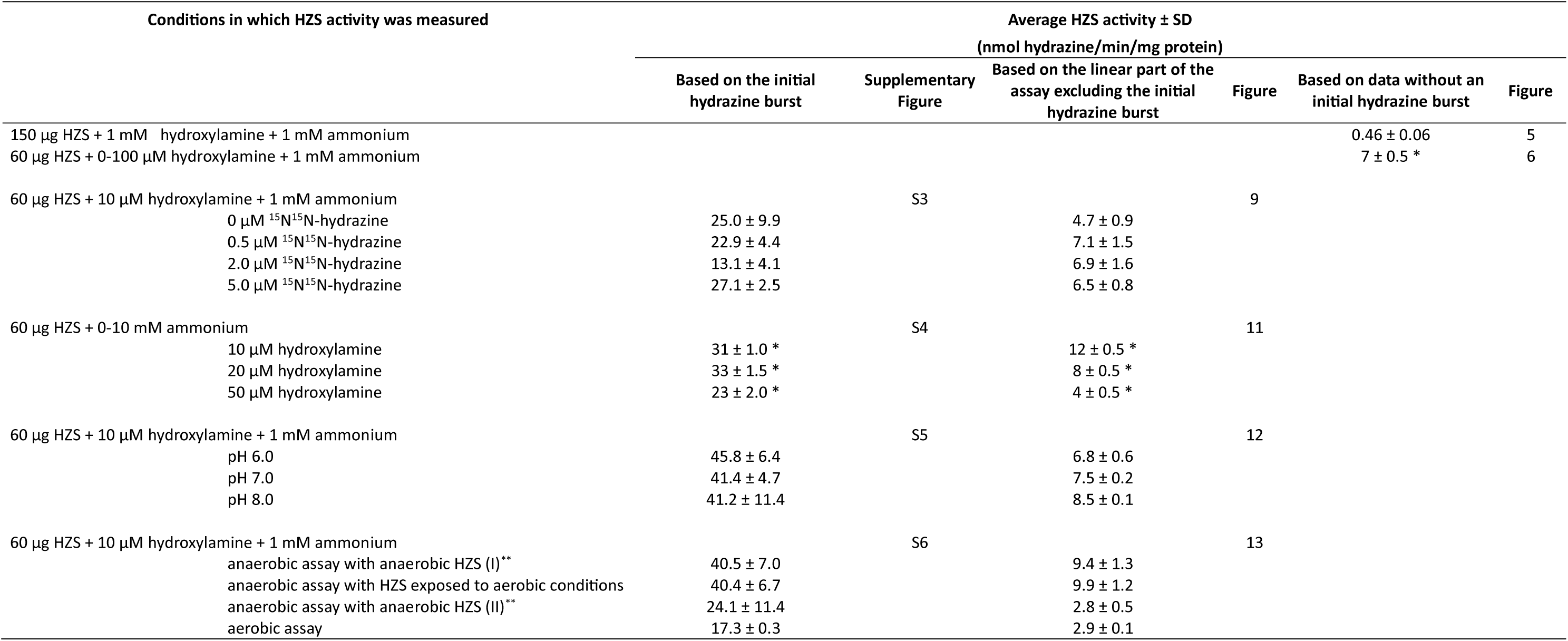
– Overview of hydrazine synthase (HZS) activity measured in different experiments via the direct enzyme assay. * Indicates *V_max_*. ** Roman number indicates a separate assay.

### Kinetic parameters of hydrazine synthase: hydroxylamine and ammonium

To determine kinetic parameters of hydrazine synthase, enzyme activity was assayed in various combinations of hydroxylamine and ammonium concentrations. Hydrazine synthase activity exhibited variability across technical replicates. Therefore, we would like to discuss the challenges in the hydrazine synthase assay, before diving into the kinetic parameters. Please note that the *K_m_* and *V_max_* values are presented as apparent, and the numbers have been rounded to reflect their indicative nature.

### Challenges in assays measuring kinetic parameters of hydrazine synthase for ammonium Initial burst of hydrazine production

In the assays measuring the kinetic parameters of hydrazine synthase for ammonium, an initial burst of hydrazine production appeared that was not observed in the assays to measure affinity of hydrazine synthase for hydroxylamine (Figure 7). The initial burst in hydrazine production was not due to residual bound hydrazine, as hydrazine was not measured in a control sample containing hydrazine synthase and substrates before starting the assay. Assay experiments with varying hydrazine synthase concentrations showed that the height of the initial burst depended on enzyme concentration, confirming that hydrazine was enzymatically produced during this period (Figure 8). Turnover calculations based on kinetics assays with hydroxylamine (Figure 6), indicated that hydrazine synthase requires 50 seconds to convert 1 mole of hydroxylamine and 1 mole of ammonium to 1 mole of hydrazine. Because the initial hydrazine burst ends within one minute, it might reflect the pre-steady state of the assay (see below). With the assay setup used, it was not possible to measure more time points at the start of the assay (*i.e.* <20 seconds after adding hydrazine synthase to substrates ammonium and hydroxylamine) to further investigate the initial hydrazine burst.

**Figure 6.**
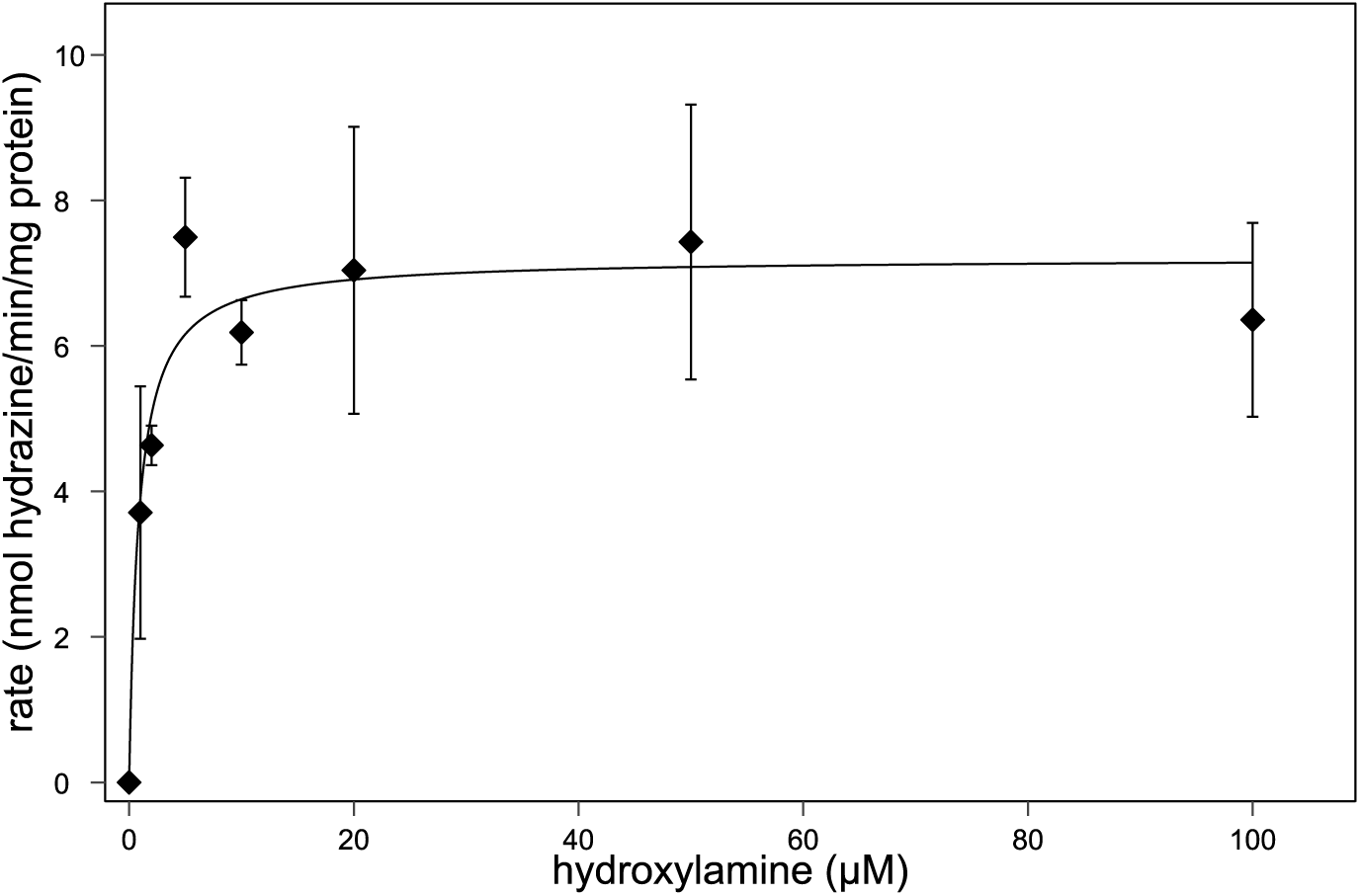
– The Michaelis-Menten parameters (*K_m_* and *V_max_*) of hydrazine synthase were determined across a range of hydroxylamine concentrations. To determine kinetic parameters of hydrazine synthase, enzyme activity was assayed at various combinations of hydroxylamine concentrations supplemented with 1 mM ammonium. The apparent *K_m_* for hydroxylamine was 1 ± 0.5 µM and the apparent *V_max_* 7 ± 0.5 nmol hydrazine/min/mg protein. Data are presented as mean ± SD (*n*=3 technical replicates).

**Figure 7.**
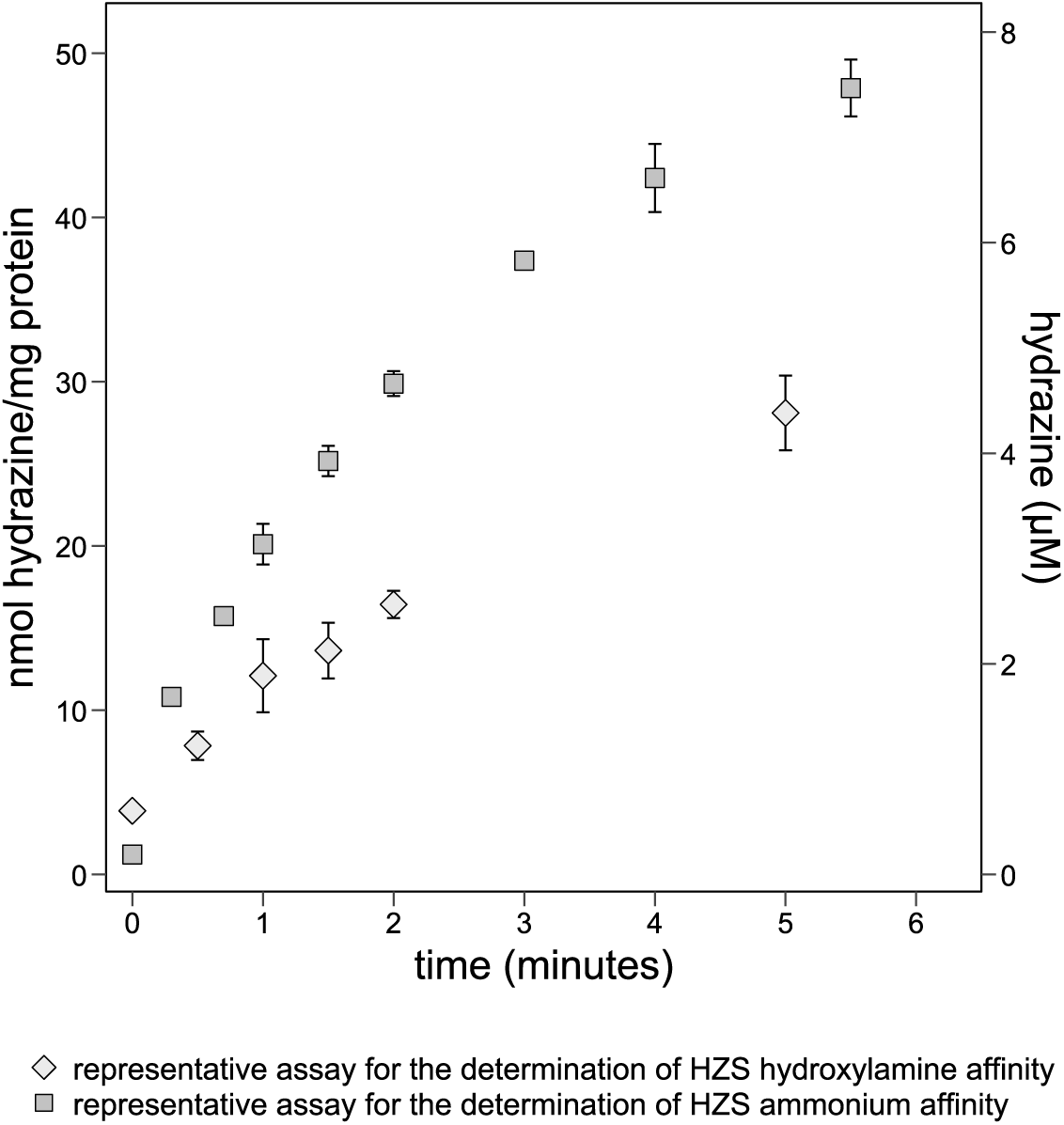
– Representative assays for the determination of hydrazine synthase ammonium and hydroxylamine affinity. In the assays to measure affinity of hydrazine synthase for ammonium, an initial burst of hydrazine production appeared (first two data points or first 30 seconds) that was not observed in the assays to measure affinity of hydrazine synthase for hydroxylamine. Activity assays to determine hydroxylamine affinity contained 60 µg hydrazine synthase, 10 µM hydroxylamine and 1 mM ammonium and activity assays to determine ammonium affinity contained 60 µg hydrazine synthase, 10 µM hydroxylamine and 10 mM ammonium. All assays were carried out in 20 mM potassium phosphate buffer, pH 7.0. Data are presented as mean ± SD (for hydroxylamine affinity assays *n*=2 technical replicates and for ammonium affinity assays *n*=3 technical replicates).

**Figure 8.**
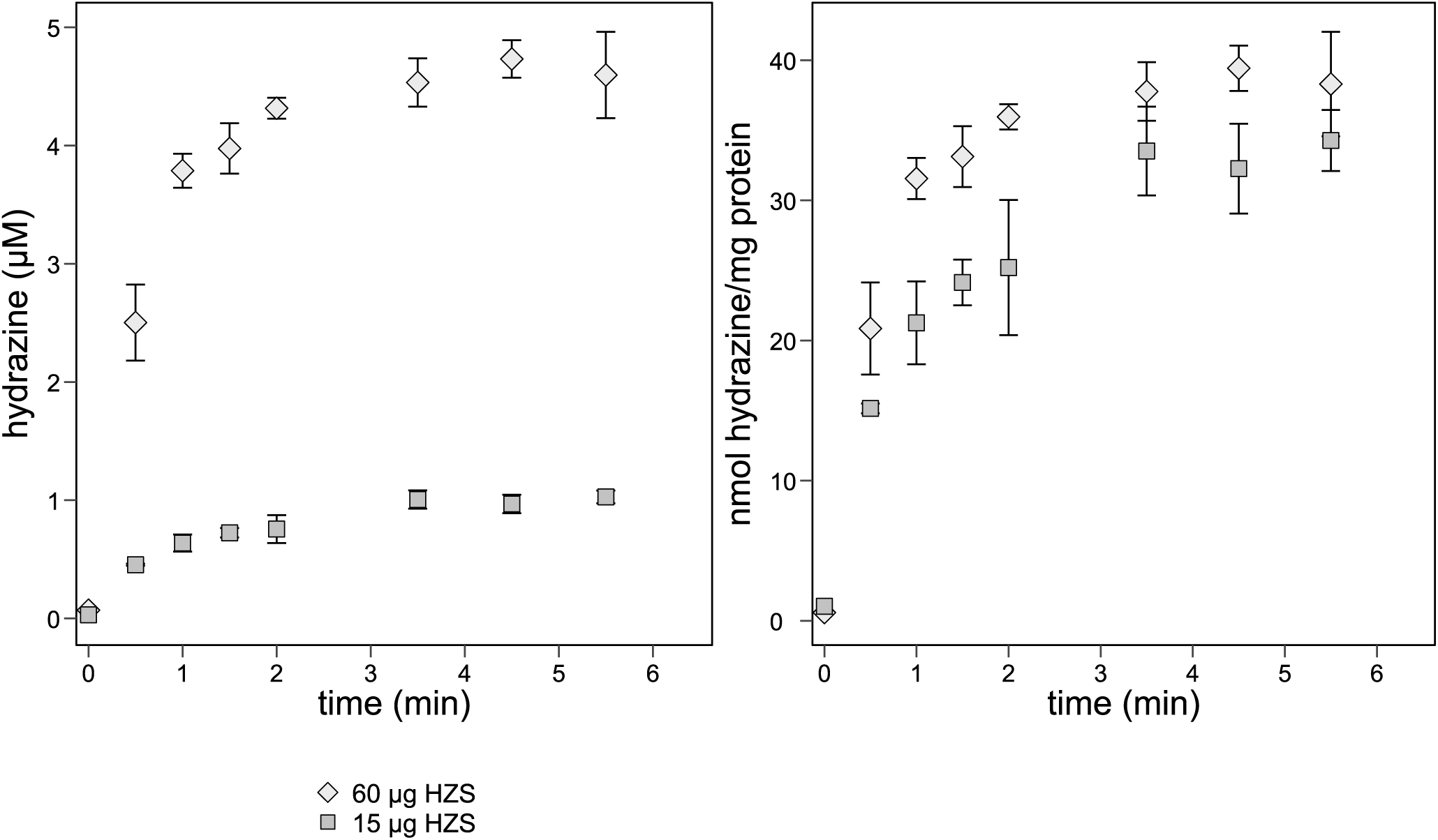
– Hydrazine synthesis measured with 60 and 15 µg hydrazine synthase. The hydrazine concentration in the initial phase of the activity assay was approximately five times lower when 15 µg hydrazine synthase was added compared to 60 µg hydrazine. The specific hydrazine synthesis in nmol/mg protein was similar for the two hydrazine synthase concentrations. Thus, hydrazine measured in the first 30 seconds of the activity assays is enzymatically produced. Activity assays contained 10 µM hydroxylamine and 1 mM ammonium and were carried out in 20 mM potassium phosphate buffer, pH 7.0. Data are presented as mean ± SD (*n*=3 technical replicates).

### Inactivation of hydrazine synthase during the activity assay

The hydrazine production seemed to decrease slightly (Figure 7) or more pronounced (Figure 8) after three to five minutes while not all hydroxylamine and ammonium were converted to hydrazine. To test whether the decreased hydrazine synthesis was caused by product inhibition of hydrazine synthase, ^15^N^15^N-hydrazine was added to the reaction mixture. ^15^N^15^N-hydrazine can be distinguished from unlabeled, biologically produced, hydrazine by LC-MS but is expected to affect hydrazine synthase in a similar way as unlabeled hydrazine. This way, the enzymatic, unlabeled hydrazine synthesis can be followed over time in the presence of ^15^N^15^N-hydrazine. Hydrazine synthesis was similar with the different ^15^N^15^N-hydrazine concentrations in the reaction mixture, ranging from 7.1 ± 1.5 to 6.5 ± 0.8 nmol hydrazine/min/mg protein for 0.5 and 5.0 µM ^15^N^15^N-hydrazine, respectively (*p*=0.8, *n*=3 technical replicates) (Figure 9, Table 2). Without additional ^15^N^15^N-hydrazine, hydrazine was produced at 4.7 ± 0.9 nmol/min/mg protein. Although this rate is lower than for the incubations with ^15^N^15^N-hydrazine, they did not differ significantly (*p*=0.1*, n*=3 technical replicates). Thus, hydrazine synthase is not inhibited by product formation below 5 µM.

**Figure 9.**
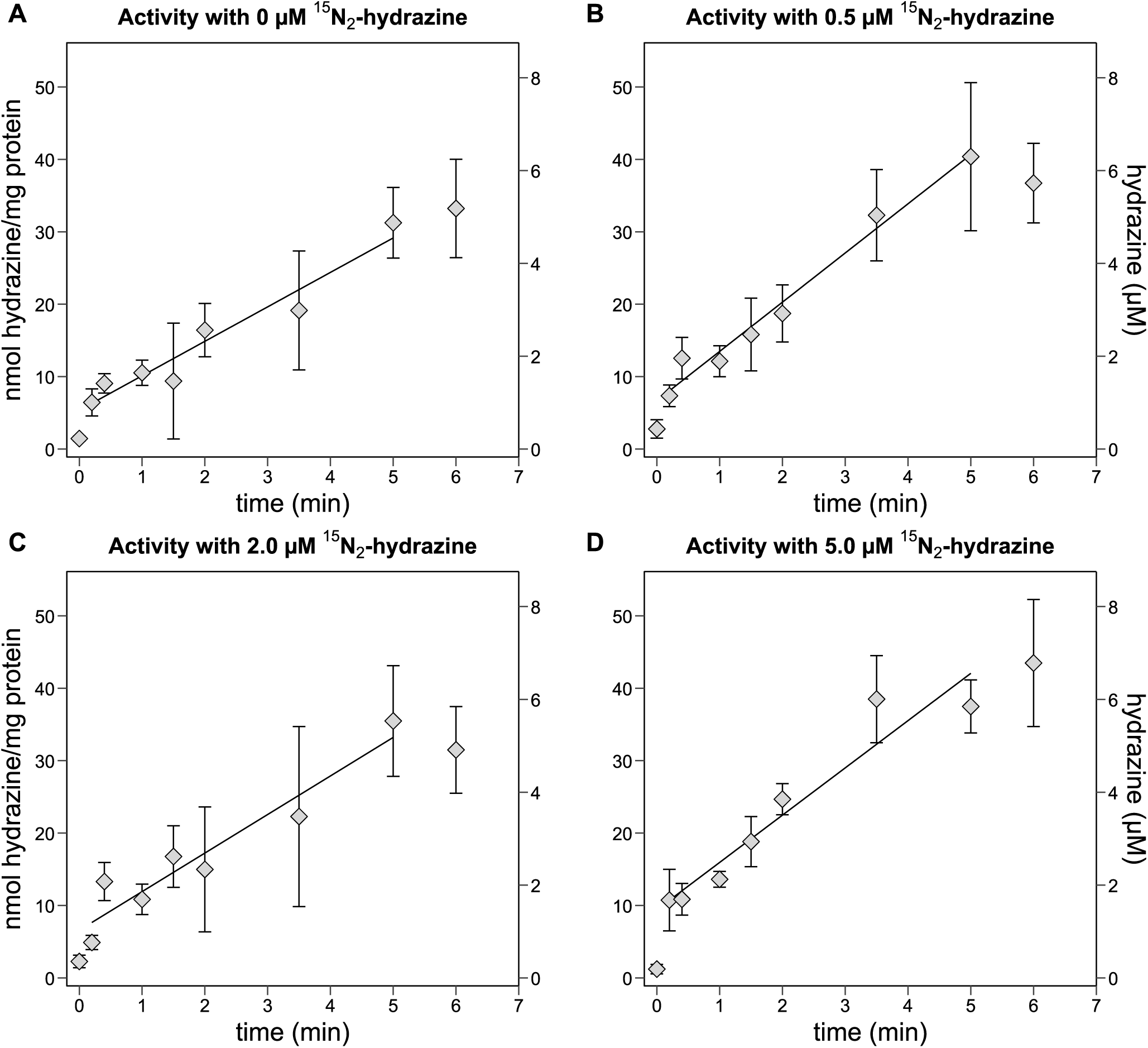
– Hydrazine synthase activity measured with various hydrazine concentrations. To measure whether hydrazine synthase was inhibited by end-product formation, activity assays (A) without ^15^N^15^N-hydrazine, (B) 0.5 µM, (C) 2.0 µM or (D) 5.0 µM ^15^N^15^N-hydrazine added to the reaction mixture were performed. There was a slight increase in the synthesis of hydrazine with increasing ^15^N^15^N-hydrazine concentration. However, the differences between incubations were not significant. Thus, hydrazine synthase is not inhibited by hydrazine formation below 5 µM. For clarity the y-axis titles were placed on the outside of the graphs. The left y-axis represents nmol hydrazine/mg protein and the right y-axis represents hydrazine (µM). Activity assays contained 60 µg hydrazine synthase with 10 µM hydroxylamine and 1 mM ammonium and were carried out in 20 mM potassium phosphate buffer, pH 7.0. Data points included to calculate the hydrazine synthesis rate are connected by a trendline. Data are presented as mean ± SD (*n*=3 technical replicates).

When calculating the rates based on the initial hydrazine burst, hydrazine synthase activity was nearly the same with varying ^15^N^15^N-hydrazine concentrations, except for the activity measured with 2.0 µM ^15^N^15^N-hydrazine. Without ^15^N^15^N-hydrazine, hydrazine synthase produced 25.0 ± 9.9 nmol hydrazine/min/mg protein. For 0.5, 2.0, and 5.0 µM ^15^N^15^N-hydrazine the rate was 22.9 ± 4.4, 13.1 ± 4.1, and 27.1 ± 2.5 nmol hydrazine/min/mg protein, respectively (Supplementary figure 3, Table 2). Why hydrazine synthase activity dropped with 2.0 µM ^15^N^15^N-hydrazine is unclear. Overall, the variation in activity does not clearly correlate with added hydrazine concentrations, suggesting that the decreased hydrazine synthase activity is unlikely to be caused by product formation up to 5 µM.

Apart from product inhibition, hydrazine synthesis could also decrease due to hydrazine synthase inactivation during the activity assay, for instance through degradation. When additional hydrazine synthase was introduced at six minutes, hydrazine was again produced (Figure 10), thus indicating that hydrazine synthase was indeed inactivated.

**Figure 10.**
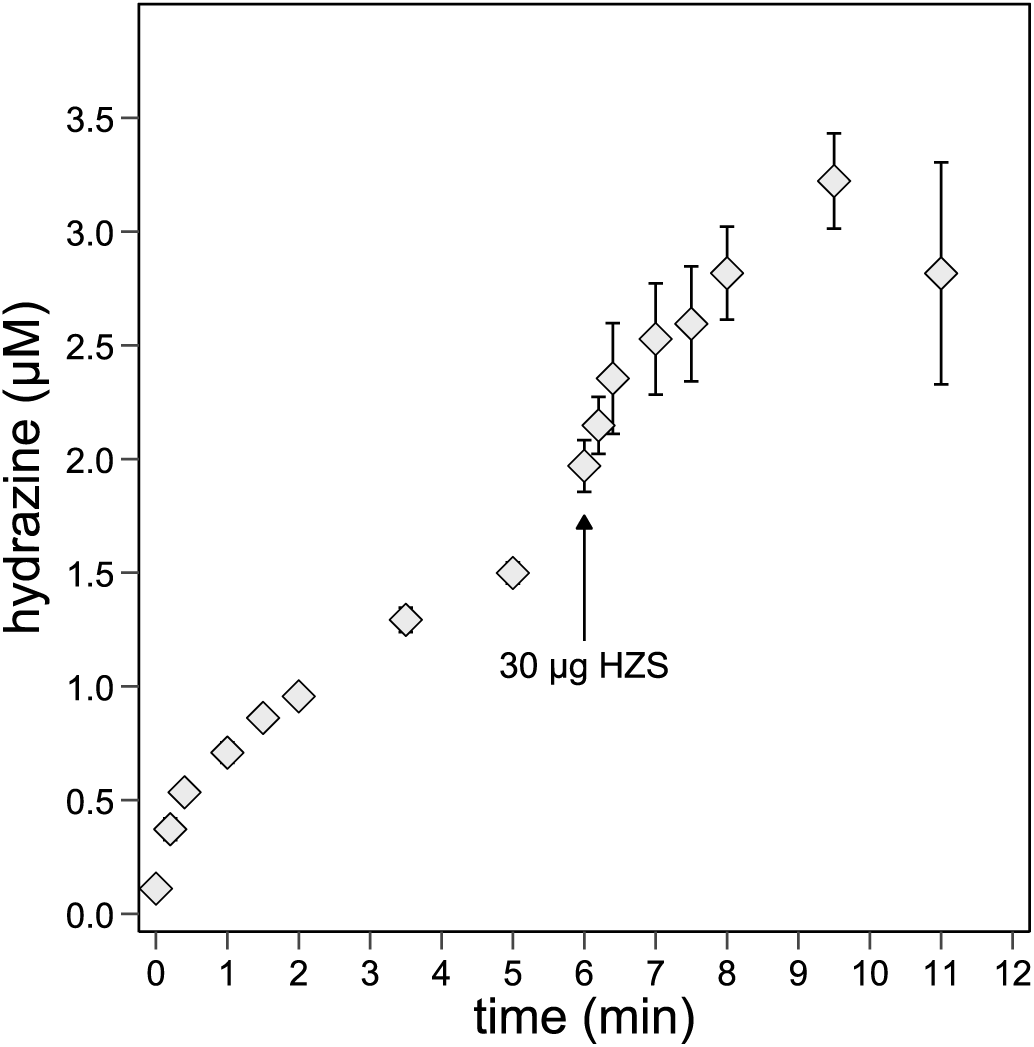
– Hydrazine synthesis measured with an addition of extra hydrazine synthase during the activity assay. To evaluate whether decreased hydrazine synthesis following the initial burst observed in assays measuring hydrazine synthase affinity for ammonium was caused by hydrazine synthase inactivation, extra enzyme was added to the reaction mixture at six minutes. Hydrazine synthesis increased after extra hydrazine synthase was added, indicating enzyme inactivation during the assay. Activity assays contained 10 µM hydroxylamine and 1 mM ammonium and were carried out in 20 mM potassium phosphate buffer, pH 7.0. Data are presented as mean ± SD (*n*=3 technical replicates).

In conclusion, the factors influencing the height of the initial hydrazine burst and the extent of hydrazine synthase inactivation remain unclear. When presenting our data, we show the calculated hydrazine synthesis rates excluding the initial burst of hydrazine production, focusing on the linear phase of the reaction with a slower synthesis rate. These results will be further analyzed in the discussion section. Nevertheless, hydrazine synthase may become inactive during the activity assay. In this case, excluding the initial hydrazine burst leads to an underestimation of the hydrazine synthesis rate in the assays. To compare the rates with and without the initial hydrazine burst, we provide the rates calculated with the initial burst in table 2 and the supplementary material.

After these considerations, let us return to the kinetic parameters *K_m_* and *V_max_* of hydrazine synthase. The apparent *K_m_* for hydroxylamine was determined at 1 ± 0.5 µM hydroxylamine with a *V_max_* of 7 ± 0.5 nmol hydrazine/min/mg protein (Table 2). A hydroxylamine concentration of 10 µM was sufficient to achieve this maximum rate (Figure 6). Moreover, when incubated with 1 mM hydroxylamine, the rate dropped to 2.0 ± 0.1 nmol hydrazine/min/mg protein which might indicate substrate inhibition effects or potential damaging of hydrazine synthase by hydroxylamine (Supplementary figure 2). These results show that hydrazine synthase has a high affinity for hydroxylamine. The turnover number (*K_cat_*) for hydrazine synthase with hydroxylamine and ammonium as substrates was determined to be 0.02/sec, indicating that hydrazine synthase is a relatively slow enzyme under our *in vitro* conditions.

The influence of hydroxylamine on hydrazine synthase was difficult to determine due to the variations between assays and the sharp bend in the Michaelis-Menten curve. Therefore, the kinetic parameters of hydrazine synthase with respect to ammonium were surveyed over a range of different hydroxylamine concentrations: 10, 20 and 50 µM.

The kinetic parameters *K_m_* and *V_max_* of hydrazine synthase with respect to ammonium were measured over a range of hydroxylamine concentrations. An apparent *V_max_* of 12 ± 0.5, 8 ± 0.5, and 4 ± 0.5 nmol hydrazine/min/mg protein was determined for 10, 20, and 50 µM hydroxylamine, respectively (Figure 11, Table 2). The apparent *K_m_* for ammonium at different hydroxylamine concentrations was 544 ± 37, 375 ± 51, and 975 ± 231 µM ammonium at 10, 20, and 50 µM hydroxylamine, respectively. More detailed analyses of the hydrazine synthase activity at varying ammonium concentrations between 10 and 800 µM with a stable hydroxylamine concentration of either 20 or 50 µM were inconclusive due to large variations in hydrazine synthesis rates (Supplementary table 1). Taken together, the data indicate that hydrazine synthase has a low affinity for ammonium.

**Figure 11.**
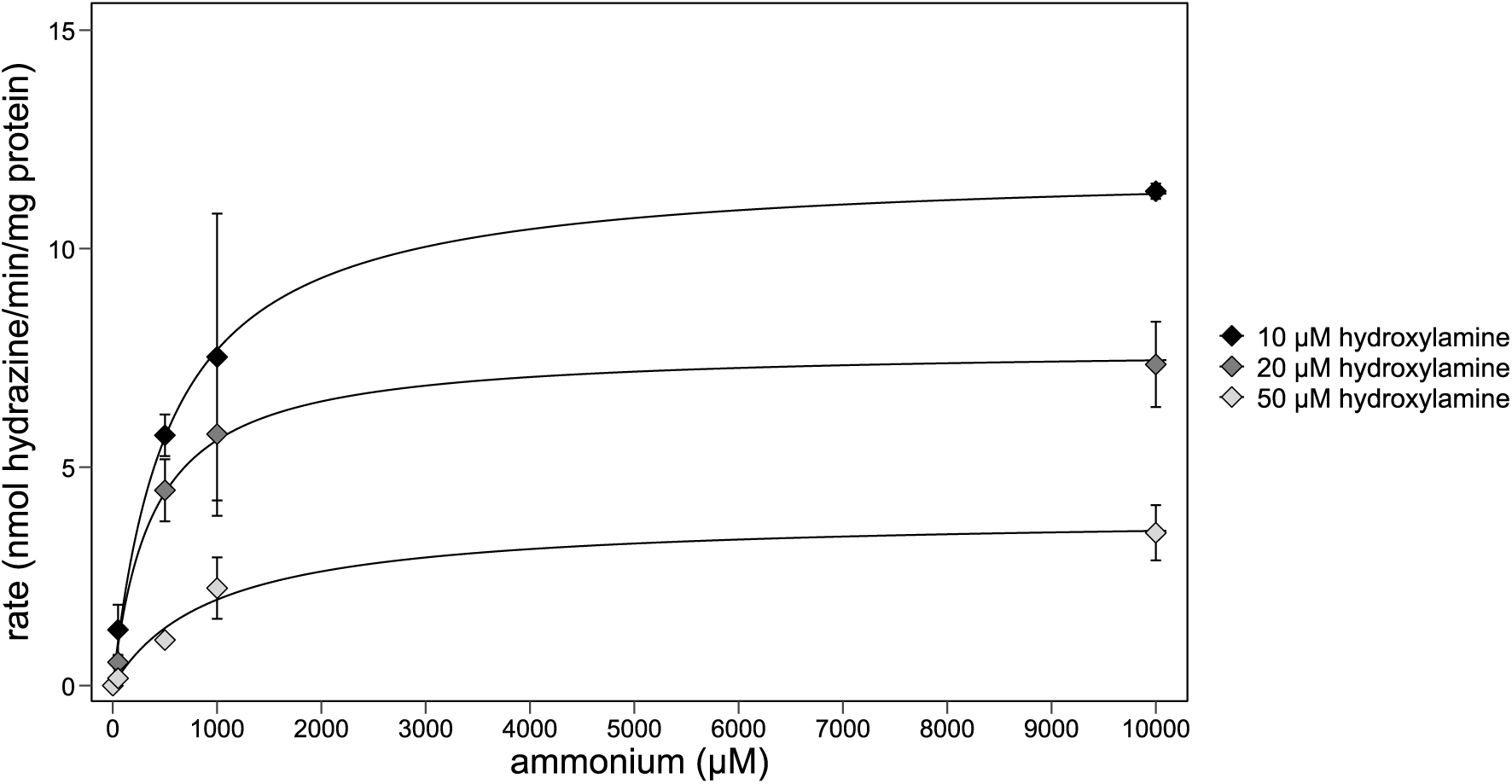
– The Michaelis-Menten parameters (*K_m_* and *V_max_*) of hydrazine synthase were determined across a range of hydroxylamine and ammonium concentrations. From incubations with 10 µM hydroxylamine the apparent *K_m_* for ammonium was determined at 544 ± 37 µM with an apparent *V_max_* of 12 ± 0.5 nmol hydrazine/min/mg protein. Incubations with 20 µM hydroxylamine showed an apparent *K_m_* of 375 ± 51 µM with an apparent *V_max_* of 8 ± 0.5 nmol hydrazine/min/mg protein. Incubations with 50 µM hydroxylamine showed an apparent *K_m_* of 975 ± 231 µM with an apparent *V_max_*of 4 ± 0.5 nmol hydrazine/min/mg protein. Please mind that for the 50 µM hydroxylamine series, the 1000 µM ammonium point is extrapolated, not measured. Data are presented as mean ± SD (*n*=3 technical replicates except for data points representing incubations with of 10 µM hydroxylamine with 1 mM ammonium, 10 µM hydroxylamine with 50 µM ammonium, and 20 µM hydroxylamine with 50 µM ammonium for which *n*=2 technical replicates).

Notably, already from 20 µM hydroxylamine inhibitory effects were observed, whereas this was only observed from 100 µM hydroxylamine in the hydroxylamine affinity experiments.

When the initial hydrazine burst, excluding the other time points, was used to determine *V_max_* and *K_m_*, hydrazine synthase seemed to be inhibited from 50 µM hydroxylamine (Supplementary figure 4, Table 2). Apparent *V_max_* was 31 ± 1.0, 33 ± 1.5, and 23 ± 2.0 nmol hydrazine/min/mg protein with an apparent *K_m_* of 75 ± 12, 107 ± 28, and 133 ± 64 µM ammonium with 10, 20, and 50 µM hydroxylamine, respectively. Thus, the datapoints included in the hydrazine synthesis rates influence the *K_m_* and *V_max_*. Nevertheless, with or without the initial hydrazine burst, hydrazine synthase likely has a low affinity for ammonium.

### Hydrazine synthase is more active at pH 8.0 than pH 6.0

To evaluate the effect of pH on hydrazine synthesis, activity was measured across three different pH values. Higher pH led to a significantly faster apparent rate of hydrazine synthesis: 6.8 ± 0.6 at pH 6.0, 7.5 ± 0.2 at pH 7.0, and 8.5 ± 0.1 nmol hydrazine/min/mg protein at pH 8.0 (*n*=3 technical replicates, Table 2). This effect may be caused directly through pH increase or indirectly by raising ammonia concentrations. To assess the potential influence of ammonia on the increased hydrazine synthesis rate, the ammonium concentration per pH was converted to the ammonia concentration using the equation: µM ammonia = (1 / (10^pka-pH^ + 1)) * µM ammonium (Emerson et al., 1975). This resulted in an ammonia range of 0.29 to 28.58 µM from pH 6.0 to 8.0, respectively. With a 100-fold increase in ammonia concentration, the reaction rate increased only 20%, suggesting that hydrazine synthesis is not strongly influenced by ammonia concentration. However, determining *V_max_*and *K_m_* is necessary to select appropriate substrate concentrations and verify whether ammonia concentration has no significant effect on hydrazine synthase activity – if already at *V_max_*, increasing the ammonia concentration would have no effect. In conclusion, the observed increase in the hydrazine synthase activity is probably primarily driven by the effect of pH on the enzyme rather than by ammonia levels.

When calculating the rates based on the initial hydrazine burst, an increasing pH did not have a stimulating effect on the hydrazine synthase activity rate. The hydrazine synthesis rates measured in pH 6.0, 7.0 and 8.0 were similar: 45.8 ± 6.4, 41.4 ± 4.7, and 41.2 ± 11.4 nmol hydrazine/min/mg protein, respectively (*n*=3 technical replicates, Table 2). Thus, the pH-range from 6.0 to 8.0 had no significant effect on hydrazine synthase activity in the burst phase.

### Hydrazine synthesis from ammonium and hydroxylamine is unaffected by oxygen

We observed that oxygen exposure during soluble protein sample preparation influenced hydrazine production from nitric oxide and ammonium, measured via dinitrogen gas in the anaerobic soluble protein fraction (Figure 2). To better understand how oxygen could impact hydrazine synthase, its effect on the second half-reaction of hydrazine production was investigated. First, anaerobically isolated hydrazine synthase was exposed to ambient air (aerobic conditions) before an anaerobic assay was performed. Untreated hydrazine synthase produced 9.4 ± 1.3 nmol hydrazine/min/mg protein and hydrazine synthase exposed to oxygen via ambient air produced 9.9 ± 1.2 nmol hydrazine/min/mg protein (Figure 12A). Secondly, activity of anaerobically isolated hydrazine synthase was examined under anaerobic and complete aerobic assay conditions (Figure 12B). Hydrazine synthase produced 2.8 ± 0.5 nmol hydrazine/min/mg protein in anaerobic conditions. In aerobic conditions, hydrazine synthase showed similar activity and produced 2.9 ± 0.1 nmol hydrazine/min/mg protein. Thus, hydrazine synthesis from hydroxylamine and ammonium by anaerobically isolated hydrazine synthase was unaffected by oxygen, whether hydrazine synthase was exposed to air before an anaerobic assay or during an aerobic assay. This might indicate that the activity loss of hydrazine synthase mainly happens in the first half-reaction where nitric oxide is reduced to hydroxylamine (Table 2).

**Figure 12.**
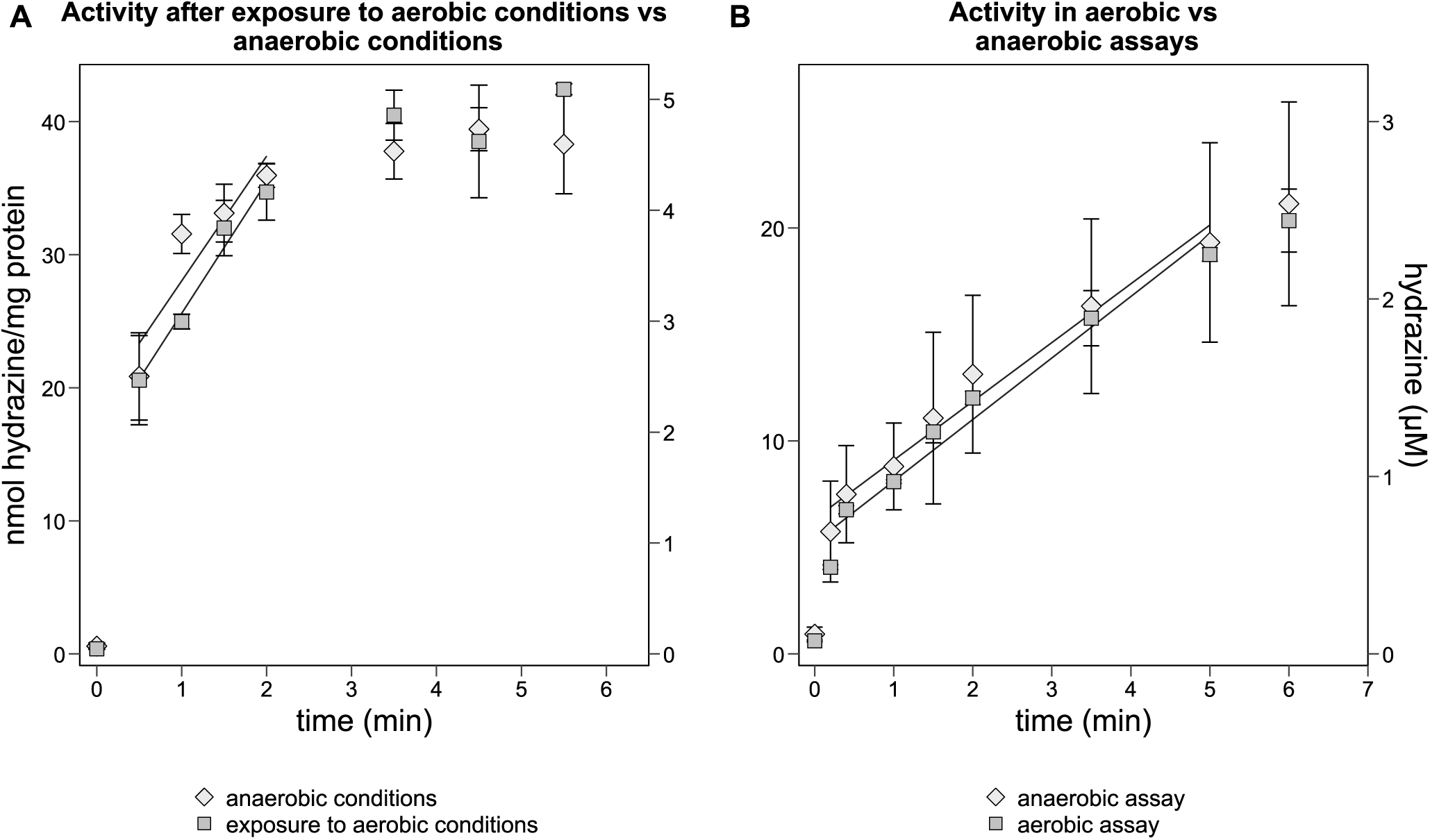
– Hydrazine synthase activity from hydroxylamine and ammonium measured in anaerobic and in aerobic conditions, and after exposure of hydrazine synthase to aerobic conditions. (A) Hydrazine synthase exposed to oxygen via ambient air produced hydrazine at a similar rate as hydrazine synthase kept in anaerobic conditions. (B) Hydrazine synthase in an aerobic assay produced hydrazine at a similar rate as hydrazine synthase in the anaerobic assay. Thus, hydrazine synthase activity from hydroxylamine and ammonium is unaffected by oxygen. Activity assays contained 60 µg hydrazine synthase with 10 µM hydroxylamine and 1 mM ammonium and were carried out in 20 mM potassium phosphate buffer pH 7.0. Data points included to calculate the hydrazine synthesis rate are connected by a trendline. Data are presented as mean ± SD (*n*=3 technical replicates).

When calculating the rates based on the initial hydrazine burst, hydrazine synthase kept under anaerobic conditions exhibited the same activity as hydrazine synthase exposed to aerobic conditions prior to an anaerobic activity assay: 40.5 ± 7.0 and 40.4 ± 6.7 nmol hydrazine/min/mg protein, respectively (Supplementary figure 5A). However, when performing the activity assay in aerobic conditions, hydrazine synthase activity was lower than in anaerobic conditions: 17.3 ± 0.3 and 24.1 ± 11.4 nmol hydrazine/min/mg protein (Supplementary figure 5B). Notably, the variation in the initial hydrazine burst between technical replicates of the anaerobic assays was rather large which influenced the conclusion (Table 2).

## Discussion

We investigated the molecular mechanism, and optimized and characterized the activity, of *K. stuttgartiensis* strain MBR1 hydrazine synthase via enzyme activity assays. We showed that oxygen negatively affects the overall hydrazine synthesis pathway from nitric oxide and ammonium in a coupled enzyme assay, but not hydrazine synthesis from hydroxylamine and ammonium in a direct assay. With the direct assay, we experimentally verified the second half-reaction of the proposed two-step hydrazine synthesis process. However, hydrazine synthase activity showed variation across assays for which the underlying mechanisms are still unclear. Despite these variations, we could determine that hydrazine synthase exhibits an apparent ammonium affinity of *K_m_* ≥ 375 ± 51 µM whereas its affinity for hydroxylamine is *K_m_* 1 ± 0.5 µM. Additionally, the hydrazine synthesis rate remains unaffected by product formation at concentrations up to 5 µM. At pH 8.0, hydrazine synthase is more active compared to pH 6.0.

### The effect of oxygen and the coupled enzyme assay on hydrazine synthesis rates

We observed that oxygen affected hydrazine production from nitric oxide and ammonium, as measured by dinitrogen gas production in the coupled enzyme assay in the anaerobic soluble protein fraction. Inhibition of hydrazine synthesis is congruent with the inhibitory effect of oxygen on the anammox process in whole cells (Jetten et al., 1997; van de Graaf et al., 1997). Hydrazine synthase may be compromised by reactive species generated from interactions between oxygen and enzymatically produced nitric oxide, hydroxylamine, and hydrazine. If oxygen indeed has an adverse effect on hydrazine synthesis by the soluble protein fraction, it might explain part of the 99% activity loss observed by Kartal and colleagues (2011b) when they compared hydrazine production by aerobically isolated hydrazine synthase to the production in whole cells. An intense low spin signal that was observed in the EPR spectrum of aerobically purified hydrazine synthase (Dietl et al., 2015), but not in the anaerobically purified hydrazine synthase (also not when exposed to oxygen) (Versantvoort et al., 2025), might be at the basis for this inactivation.

In addition, we noticed that the availability of electron acceptors and donors substantially influenced hydrazine production rates in the MIMS coupled enzyme assay. Initially, we measured a hydrazine synthesis rate of 0.03 nmol/min/mg protein. This was ten times lower than reported by Kartal et al. (2011b) for similar assays. Adding extra oxidized cytochrome *c* increased the activity rate to 22 nmol/min/mg protein. Although, this rate may not reflect actual hydrazine production, because hydrazine is not measured directly but via dinitrogen gas production, it shows that an accurate determination of hydrazine synthesis in the coupled assay requires balancing electron donor and acceptor pools. Imbalances may further explain the discrepancy between hydrazine synthesis rates in whole cells and by isolated hydrazine synthase.

### Direct hydrazine quantification

We measured the hypothesized second half-reaction of hydrazine production via the quantification of derivatized hydrazine to obtain more insight into the molecular mechanism of hydrazine synthase. We were able to experimentally verify that isolated hydrazine synthase produces hydrazine from externally supplied hydroxylamine and ammonium. The highest apparent maximum rate obtained was 12.0 ± 0.5 nmol hydrazine/min/mg protein, which is 35 times higher than previously reported for hydrazine synthase enzyme assays with nitric oxide and ammonium to measure the overall hydrazine synthesis reaction (Kartal et al., 2011b). In addition, this activity rate was 12.5 times lower than the potential rate of hydrazine synthase calculated in whole cells. Thus, the combination of hydroxylamine and ammonium as substrates and direct quantification of hydrazine markedly improved hydrazine synthesis rates measured *in vitro*. Nevertheless, comparing results of the two different methods used to determine hydrazine synthase activity is difficult.

In most enzyme assay conditions, we observed an initial burst of hydrazine followed by a slower production rate. The cause of this phenomenon remains unclear. Enzyme turnover calculations indicated that hydrazine synthase converts hydroxylamine and ammonium to hydrazine at a relatively slow rate (*K_cat_* of 0.02/sec). This suggests that hydrazine synthesis slowed down before the enzyme could complete a single turnover, possibly indicating that the observed activity burst occurs in the pre-steady state.

We also demonstrate evidence that hydrazine synthase is inactivated during the activity assay (Figure 10). Possibly, hydroxylamine slowly inactivates hydrazine synthase. However, no clear correlation was observed between hydroxylamine concentration ≤100 µM and activity decline when determining enzyme kinetics of hydrazine synthase for hydroxylamine (Figure 6). In contrast, an inhibitory effect from 20 µM hydroxylamine was observed in enzyme kinetics experiments with ammonium and hydroxylamine (Figure 11). This effect became more pronounced with 50 µM hydroxylamine. In case hydroxylamine slowly inhibits hydrazine synthase, excluding the initial hydrazine burst leads to an underestimation of the hydrazine synthesis rate in most assays. To further investigate the initial hydrazine burst, more time-points in this part of the enzyme assay should be taken. This could be achieved by slowing hydrazine synthesis in the assay, for instance by lowering enzyme and hydroxylamine concentrations. In addition, it would be interesting to examine the effect of temperature. This was, however, challenging in the current set-up with the anaerobic glove box operating at 16°C.

### Molecular mechanism of hydrazine synthase

Our findings indicated that *K. stuttgartiensis* hydrazine synthase can utilize externally supplied hydroxylamine, with an apparent *K_m_* of 1 ± 0.5 µM and *V_max_* achieved at 10 µM hydroxylamine (Figure 6). However, due to inconsistencies in inhibitory hydroxylamine concentrations across assays—while some assays showed hydrazine synthase inhibition at 20 µM hydroxylamine (Figure 11), others only at 1 mM (Supplementary Figure 2)—we were unable to unambiguously determine the effect of hydroxylamine on hydrazine synthase.

In anammox metabolism research, the role of hydroxylamine as a free intermediate and substrate for hydrazine synthase remains a subject of ongoing debate. Studies on *K. stuttgartiensis* suggest that free hydroxylamine has no significant role in the anammox reaction. Consistent with this, Kartal et al. (2013) hypothesized that hydroxylamine leaking from hydrazine synthase is rapidly recycled by HOX into the hydrazine synthase substrate nitric oxide (Kartal et al., 2013; Maalcke et al., 2014). On the contrary, Oshiki et al. (2016a) argued that hydroxylamine is important in the anammox reaction and serves as a free intermediate and substrate for “*Candidatus* Brocadia” hydrazine synthase.

In the present study, we could neither confirm nor refute the role of hydroxylamine as free intermediate and substrate for hydrazine synthase, but indicate that, for *K. stuttgartiensis,* its presence as a free substrate might be risky. We observed hydrazine synthase inhibition *in vitro* at hydroxylamine concentrations starting from 20 µM and potentially during activity assays after two minutes. Similarly, Maalcke et al. (2016) reported inhibition of hydrazine dehydrogenase (HDH) by hydroxylamine at concentrations starting from 8 µM. HDH catalyzes the last step of the anammox reaction in which hydrazine is oxidized to dinitrogen gas (Maalcke et al., 2016). If hydroxylamine inhibits both hydrazine synthase and HDH, HOX may play an important role in removing hydroxylamine to maintain metabolic stability. However, care should be taken when translating *in vitro* results to *in vivo*. Overall, the role of hydroxylamine in anammox bacteria and its interaction with hydrazine synthase warrant further investigation.

In contrast to the high affinity for hydroxylamine, hydrazine synthase exhibits low affinity for ammonium with an apparent *K_m_* of 375 µM–1 mM. Although this *K_m_* is probably an underestimation due to assay conditions, it aligns with earlier theories on the low affinity of hydrazine synthase for ammonium formulated by Smeulders et al. (2020). When *K. stuttgartiensis* cells were grown under ammonium limited conditions (5 mM ammonium) the hydrazine synthase gene cluster was upregulated, indicating that ammonium affinity of hydrazine synthase is insufficient to maintain the rate of hydrazine synthesis at low ammonium levels (Smeulders et al., 2020). In agreement with the hypothesis that hydrazine synthase has a low affinity for ammonium, hydrazine synthase constitutes a major portion (20%) of the total protein content of *K. stuttgartiensis* cells when ammonium is abundant (45 mM) (Kartal et al., 2013), possibly to maintain sufficient hydrazine concentrations despite its low ammonium affinity.

An alternative hypothesis is that ammonia, rather than ammonium, serves as the actual substrate for hydrazine synthase. Su et al. (2021) proposed that ammonium is unlikely to directly participate in hydrazine formation due to unfavorable energetics. Instead, ammonia could react with hydroxylamine to hydrazine. To acquire ammonia, the zinc site near heme αI could remove a proton from ammonium, converting it into ammonia (Su et al., 2021). Alternatively, ammonia might enter the αI site through the minor tunnel. If ammonia serves as the substrate for hydrazine synthase, our data suggest that hydrazine synthase has a high affinity for this substrate (*K_m_* ∼1.0 ± 0.2 µM).

Furthermore, we observed that isolated hydrazine synthase is more active at pH 8.0 compared to pH 6.0. Analogously, thermodynamic calculations and experiments showed that higher pH leads to increased hydrazine production in anammox cells (Soler-Jofra et al., 2020). However, the pH in the anammoxosome, the location of the anammox energy metabolism, is proposed to be 6.3 (van der Star et al., 2010). How the pH influences the two half-reactions of hydrazine synthesis in the cell would be an interesting topic for future research.

Lastly, hydrazine synthesis from hydroxylamine and ammonium by anaerobically isolated hydrazine synthase is unaffected by oxygen. Su et al. (2021) previously proposed that hydroxylamine reacts at the Fe^II^ of catalytic heme αI to Fe^III^-NH_2_ + ·OH radical. However, in anaerobically isolated hydrazine synthase this heme αI was in the oxidized Fe^III^ state and redox titrations revealed that it has a very low redox potential below –610 mV, indicating that the reaction takes place on an oxidized heme instead (Versantvoort et al., 2025). This, together with our observation that hydrazine synthesis from hydroxylamine and ammonium is unaffected in the oxidized state of hydrazine synthase, suggests that condensation of hydroxylamine and ammonia occurs on an oxidized heme.

## Conclusion

Here, we show that oxygen affects the overall hydrazine synthesis pathway from nitric oxide and ammonium *in vitro*. This effect was minimized using an anaerobic isolation procedure. Moreover, we experimentally verified the second half-reaction in the proposed two-step mechanism for hydrazine production by hydrazine synthase. Anaerobically isolated *K. stuttgartiensis* strain MBR1 hydrazine synthase is a slow enzyme and can use externally supplied hydroxylamine and ammonium, yielding hydrazine. However, we were unable to confirm or exclude the role of hydroxylamine as a free intermediate and substrate for hydrazine synthase. Our findings suggest that hydroxylamine as a free substrate may pose a risk to *K. stuttgartiensis* energy conservation. Contradictory to the high affinity of hydrazine synthase for hydroxylamine, its affinity for ammonium is low. This may explain the abundance of the enzyme in anammox bacteria, enabling sufficient hydrazine synthesis for energy conservation. Lastly, we show that the condensation of hydroxylamine and ammonium is unaffected by oxygen, strongly suggesting that heme αI initiates this reaction in oxidized Fe^III^ state. Although activity assays are a powerful tool to study the molecular mechanisms of enzymes, they may not always accurately reflect the enzyme’s behavior *in vivo*. Moreover, activity assays with isolated hydrazine synthase, hydroxylamine and ammonium are technically and experimentally challenging but warrant further investigation to fully understand the interactions of hydrazine synthase with these substrates. Ultimately, these new insights in the molecular mechanism of hydrazine synthase bring us a step closer in unravelling the cellular metabolism of the intriguing anammox bacteria and pave the way for a biotechnological application of hydrazine synthase in sustainable and biological hydrazine production from wastewater.

## Supporting information

This article contains supporting information

## Acknowledgements

We would like to thank Guylaine Nuijten for assistance and co-maintenance of anammox bioreactor enrichment cultures, Lotte Nijman for the SDS-PAGE image in Supplementary figure 1, and Arjan Pol for helpful discussions.

## Funding

LvN, FJV and WV were supported by VIDI grant VI.Vidi.192.001 awarded to LvN by the Dutch Research Council (NWO). LvN and SCMH-V were supported by an Interdisciplinary Research Platform voucher awared to LvN from the Faculty of Science (Radboud University).

## Author contributions

LvN obtained funding and conceptualized the project, LvN, RSJ and WV supervised the project, all authors designed experiments, FJV, SCMH-V, PMMvdV, RM and WV performed experiments, all authors analyzed data, FJV and LvN wrote the first draft of the manuscript and all authors reviewed, edited and approved the manuscript.

## Conflict of interest

The authors have no conflict of interest to declare.

## Data accessibility

The data that supports the findings of this study are available within the article and in the supplementary material. In addition, biological samples used in this study are available from the corresponding author upon reasonable request.

**This article contains supporting information**.

## Notes

### Competing Interest Statement

The authors have declared no competing interest.

