## Supplementary material for "The molecular mechanism and activity of *Kuenenia stuttgartiensis* hydrazine synthase": This article contains supporting information

### Supplementary data

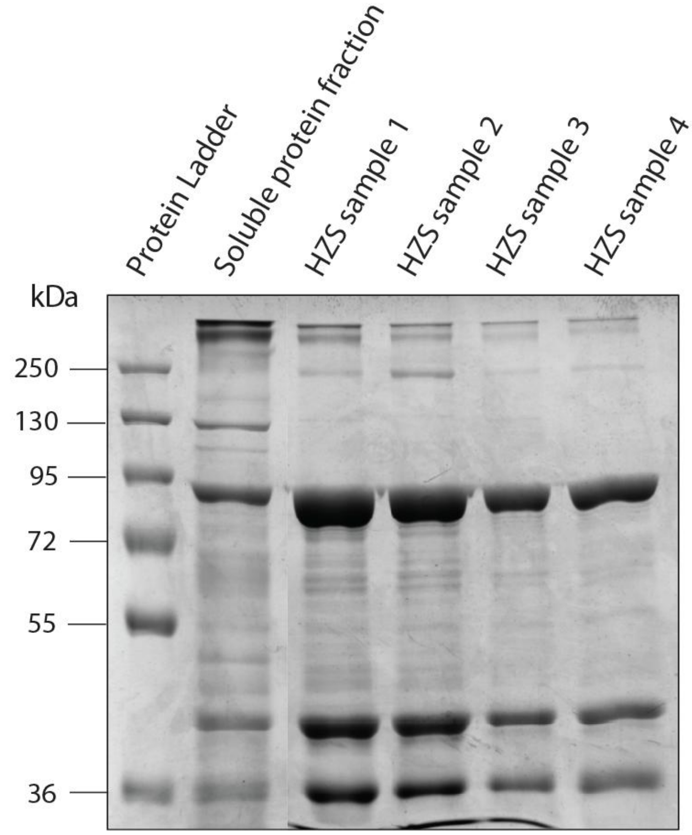


**Supplementary figure 1** – SDS-PAGE of isolated *K. stuttgartiensis* hydrazine synthase samples. Hydrazine synthase shows three bands representing subunit α (95 kDa), β (50 kDa), and γ (36 kDa). Of every sample, 15 µg protein was loaded. Photograph courtesy of L. Nijman.

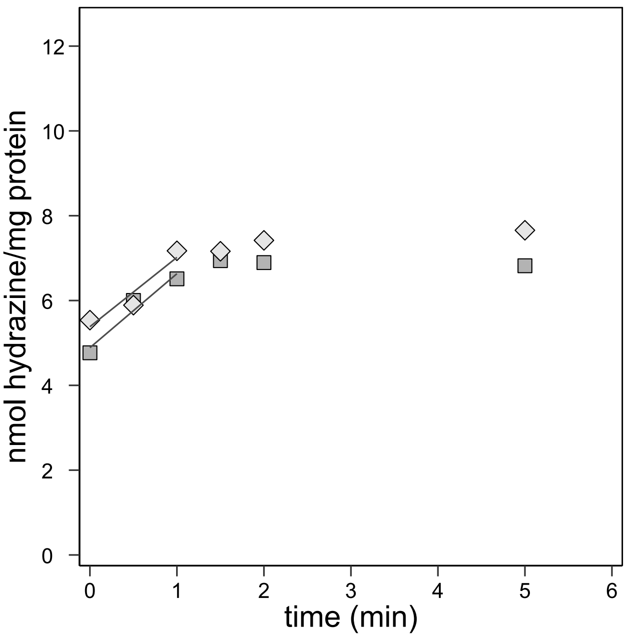


**Supplementary figure 2** – Relatively low hydrazine synthesis rate of 2.0 nmol hydrazine/min/mg protein from 1 mM hydroxylamine and 1 mM ammonium. Data points included to calculate the hydrazine synthesis rate are connected by a trendline. Every symbol represents one technical replicate.

**Supplementary table 1** – Analyses of hydrazine synthesis with ammonium concentrations ≤ 800 µM. To determine the kinetic parameters of hydrazine synthase, hydrazine synthesis was measured at various ammonium concentrations between 10 and 800 µM. The variability in hydrazine synthesis rates between the three technical replicates is represented by the large standard deviation (SD). Furthermore, the $R^2$ determined for every rate showed that the variation in hydrazine synthesis rates could not be explained by changing ammonium concentrations. Thus, the lower ammonium concentrations yielded inconclusive results on hydrazine synthase affinity for ammonium. Data are presented as mean ± SD (*n*=3 technical replicates).

| | **20 µM hydroxylamine** | |
|---|---|---|
| **Ammonium (µM)** | **rate ± SD (nmol hydrazine/min/mg protein)** | **$R^2$ ± SD** |
| 10 | 0.11 ± 0.08 | 0.62 ± 0.35 |
| 20 | 0.01 ± 0.05 | 0.29 ± 0.19 |
| 50 | 0.12 ± 0.02 | 0.76 ± 0.24 |
| 100 | 0.15 ± 0.12 | 0.60 ± 0.34 |
| 200 | 0.31 ± 0.13 | 0.90 ± 0.10 |
| 400 | 0.30 ± 0.30 | 0.60 ± 0.24 |
| 800 | 0.60 ± 0.34 | 0.90 ± 0.05 |

| | **50 µM hydroxylamine** | |
|---|---|---|
| **Ammonium (µM)** | **rate ± SD (nmol hydrazine/min/mg protein)** | **$R^2$ ± SD** |
| 1 | 0.42 ± 0.33 | 0.90 ± 0.10 |
| 2 | 0.34 ± 0.16 | 0.65 ± 0.24 |
| 5 | 0.34 ± 0.67 | 0.37 ± 0.55 |
| 10 | 0.26 ± 0.16 | 0.71 ± 0.05 |
| 20 | 1.00 ± 0.23 | 0.82 ± 0.15 |
| 50 | 1.77 ± 0.97 | 0.88 ± 0.07 |
| 100 | 0.89 ± 0.24 | 0.95 ± 0.20 |

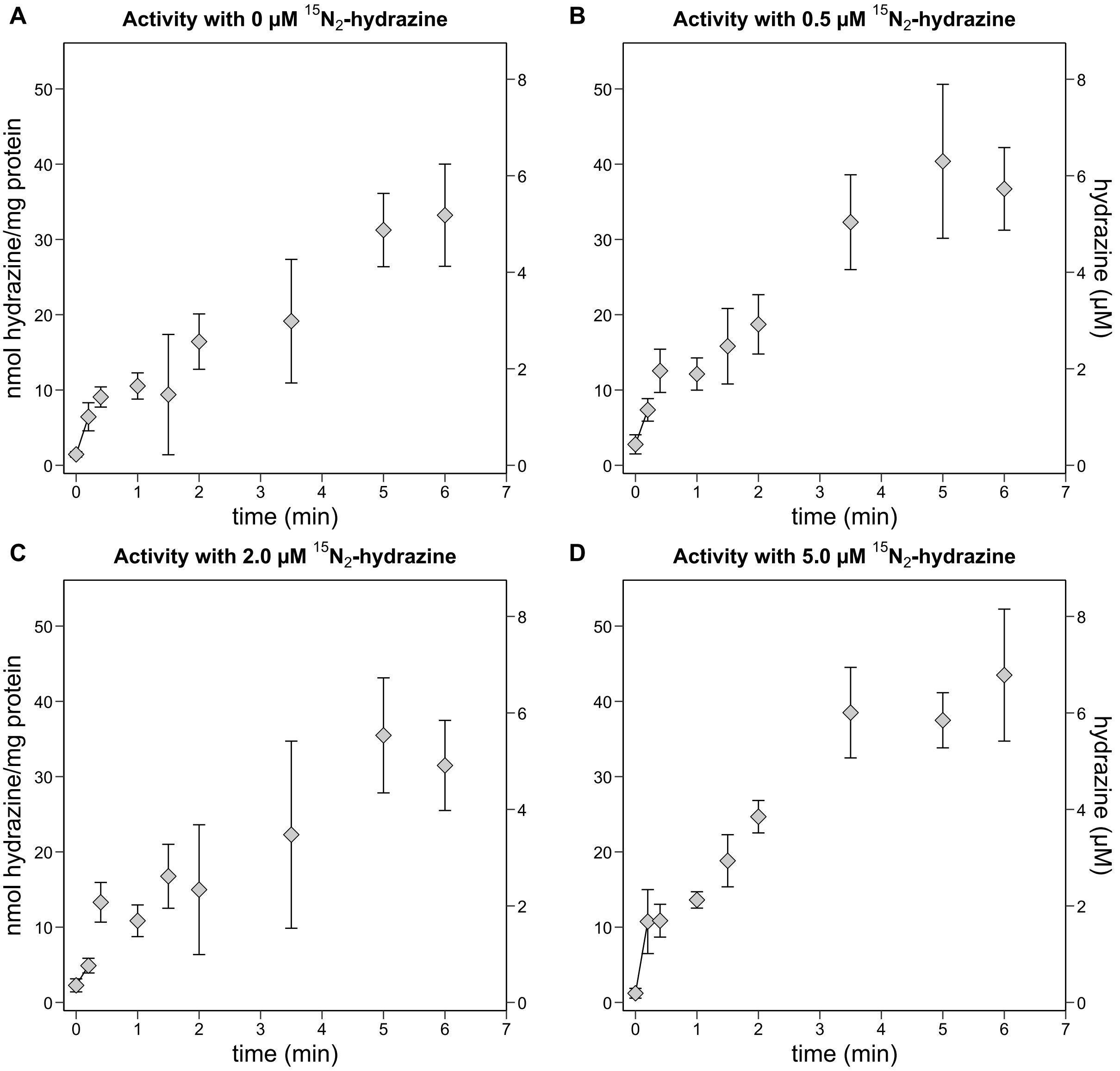


**Supplementary figure 3** – Hydrazine synthase activity measured with various hydrazine concentrations. To measure whether hydrazine synthase was inhibited by end-product formation, activity assays (A) without $^{15}N^{15}N$-hydrazine, (B) 0.5 µM, (C) 2.0 µM or (D) 5.0 µM $^{15}N^{15}N$-hydrazine added to the reaction mixture were performed. Hydrazine synthase activity is nearly similar with the varying $^{15}N^{15}N$-hydrazine concentrations, except for the activity measured with 2.0 µM $^{15}N^{15}N$-hydrazine. Thus, variation in activity does not clearly correlate with added hydrazine concentrations, suggesting that the decreased hydrazine synthase activity is unlikely to be caused by product formation up to 5 µM. For clarity the y-axis titles were placed on the outside of the graphs. The left y-axis represents nmol hydrazine/mg protein and the right y-axis represents hydrazine (µM). Activity assays contained 60 µg hydrazine synthase with 10 µM hydroxylamine and 1 mM ammonium and were carried out in 20 mM potassium phosphate buffer, pH 7.0. Data points included to calculate the hydrazine synthesis rate are connected by a trendline. Data are presented as mean ± SD ($n$=3 technical replicates).

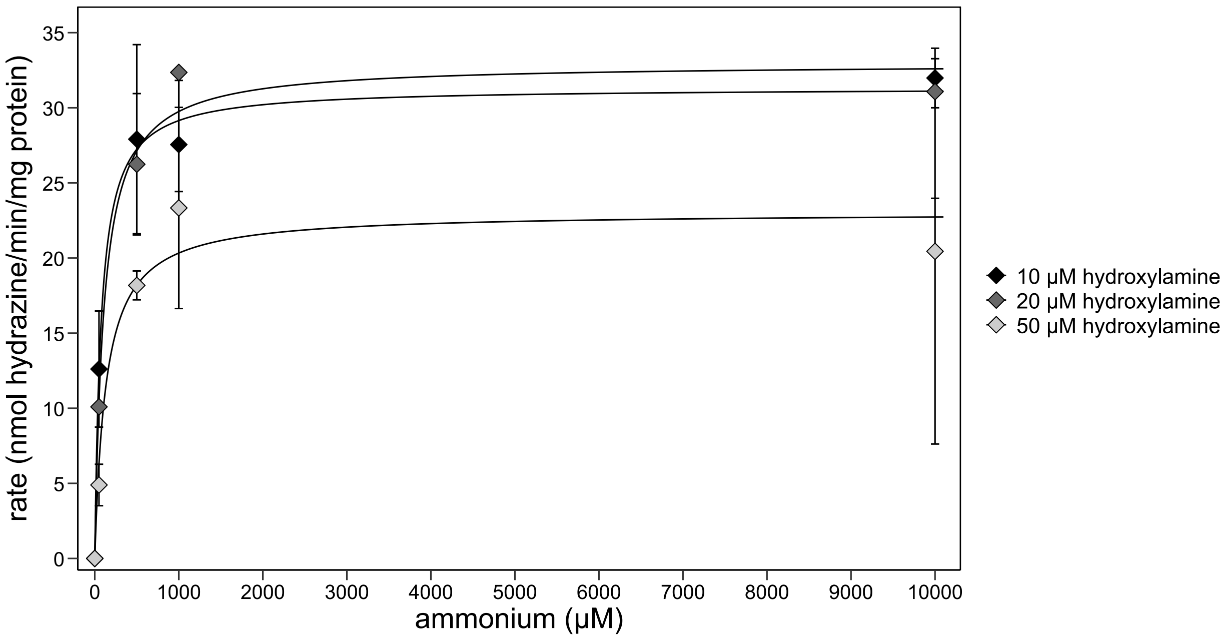


**Supplementary figure 4** – The Michaelis-Menten parameters ($K_m$ and $V_{max}$) of hydrazine synthase were determined across a range of hydroxylamine and ammonium concentrations. Hydrazine synthase has a low affinity for ammonium. For the determination of the two parameters, only the initial burst in hydrazine production was included. From incubations with 10 µM hydroxylamine the apparent $K_m$ for ammonium was determined at 75 ± 12 µM with an apparent $V_{max}$ of 31 ± 1.0 nmol hydrazine/min/mg protein. Incubations with 20 µM hydroxylamine showed an apparent $K_m$ of 107 ± 28 µM with an apparent $V_{max}$ of 33 ± 1.5 nmol hydrazine/min/mg protein. Incubations with 50 µM hydroxylamine showed an apparent $K_m$ of 133 ± 64 µM with an apparent $V_{max}$ of 23 ± 2.0 nmol hydrazine/min/mg protein. Data are presented as mean ± SD ($n$=3 technical replicates except for data points representing incubations with of 10 µM hydroxylamine with 1 mM ammonium, 10 µM hydroxylamine with 50 µM ammonium, and 20 µM hydroxylamine with 50 µM ammonium for which $n$=2 technical replicates).

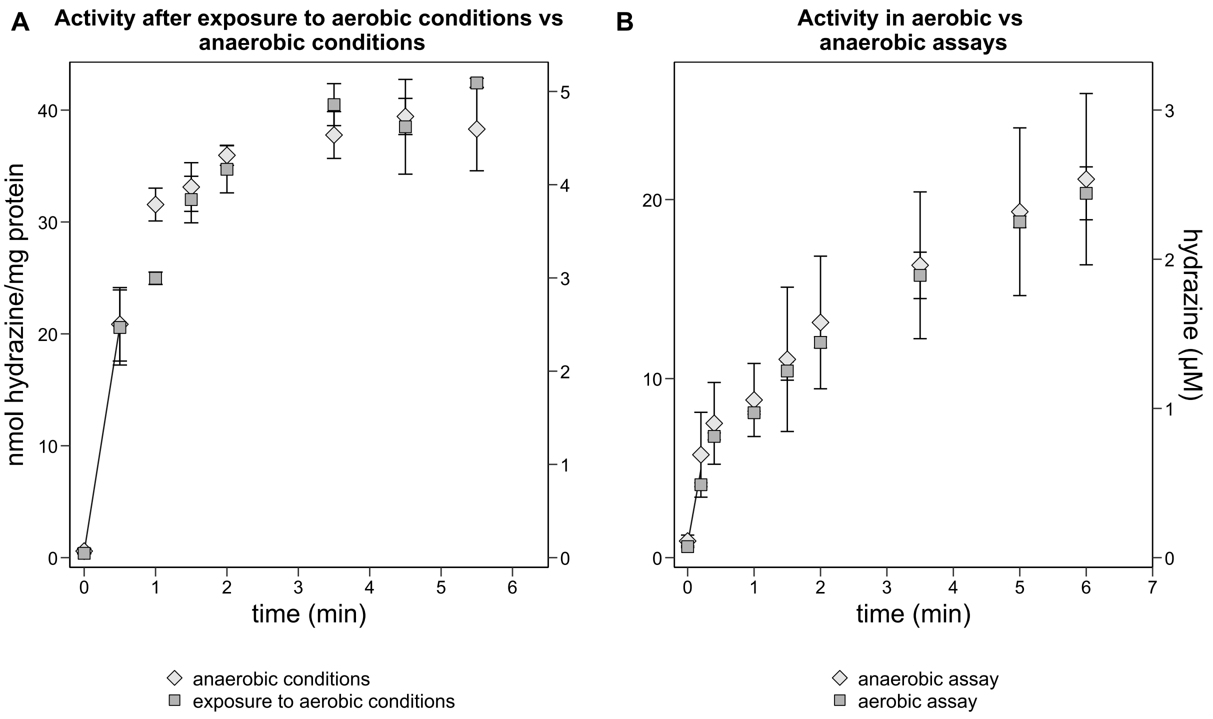


**Supplementary figure 5** – Hydrazine synthase activity from hydroxylamine and ammonium measured in anaerobic, in aerobic conditions, and after exposure of hydrazine synthase to aerobic conditions. (A) Hydrazine synthase exposed to oxygen via ambient air produced hydrazine at a similar rate as hydrazine synthase kept in anaerobic conditions. (B) Hydrazine synthase in an aerobic assay produced hydrazine at a lower rate than hydrazine synthase in the anaerobic assay. Notably, the variation in the initial hydrazine burst between technical replicates of the anaerobic assay was rather large, influencing the conclusion. Activity assays contained 60 µg hydrazine synthase with 10 µM hydroxylamine and 1 mM ammonium and were carried out in 20 mM potassium phosphate buffer pH 7.0. Data points included to calculate the hydrazine synthesis rate are connected by a trendline. Data are presented as mean ± SD ($n$=3 technical replicates).
